# A λ Phage Platform for Successful Therapeutic Display of Protein Antigens

**DOI:** 10.64898/2026.01.19.700330

**Authors:** Meredith Bush, Xintian Li, Manoj Rajaure, Donald L. Court, Sankar Adhya

**Affiliations:** Laboratory of Molecular Biology, Center for Cancer Research, National Cancer Institute, National Institutes of Health, Bethesda, MD, USA; Formerly Meredith Showler; Laboratory of RNA Biology, Center for Cancer Research, National Cancer Institute, NIH, Frederick, MD, USA

## Abstract

We have developed a vector platform for delivery of foreign peptides by genetic modification of the temperate lambda (λ) bacteriophage. This delivery platform is capable of displaying peptides or proteins on either terminus of the structural λ head protein D, present in ∼420 copies per phage particle, and λ side tail fiber (Stf), present at 12 copies per phage particle. Proteins and peptides can be easily fused for display through the low-cost and high-efficiency methods of recombineering and λ prophage induction for recombinant phage preparation described here. To improve this vector technology for use in antigen selection and immunotherapy, we introduced several mutations in the bacterial host and resident prophage λ that improve engineering, induction, phage stability, yield, fusion protein accommodation capacity, and longevity in animal systems. We tested the ability of this λ display system to identify useful antigens and generate antibodies in a mouse model. We report its success as a new technology for both applications: the selection and delivery of therapeutic peptides and proteins.

## Introduction

Bacteriophages have been used since the 1990s to display foreign peptides or proteins on the surface of their virion structures^1,2^. The bacteriophage used here as a platform for display is lambda (λ), an *Escherichia coli (E. coli)* bacteriophage composed of two major structural elements (Fig. 1A): the phage capsid or head, which contains the genome, and the phage tail, which includes proteins for binding the bacterial receptor (LamB) and triggering genome injection^3^.

**Fig 1.**
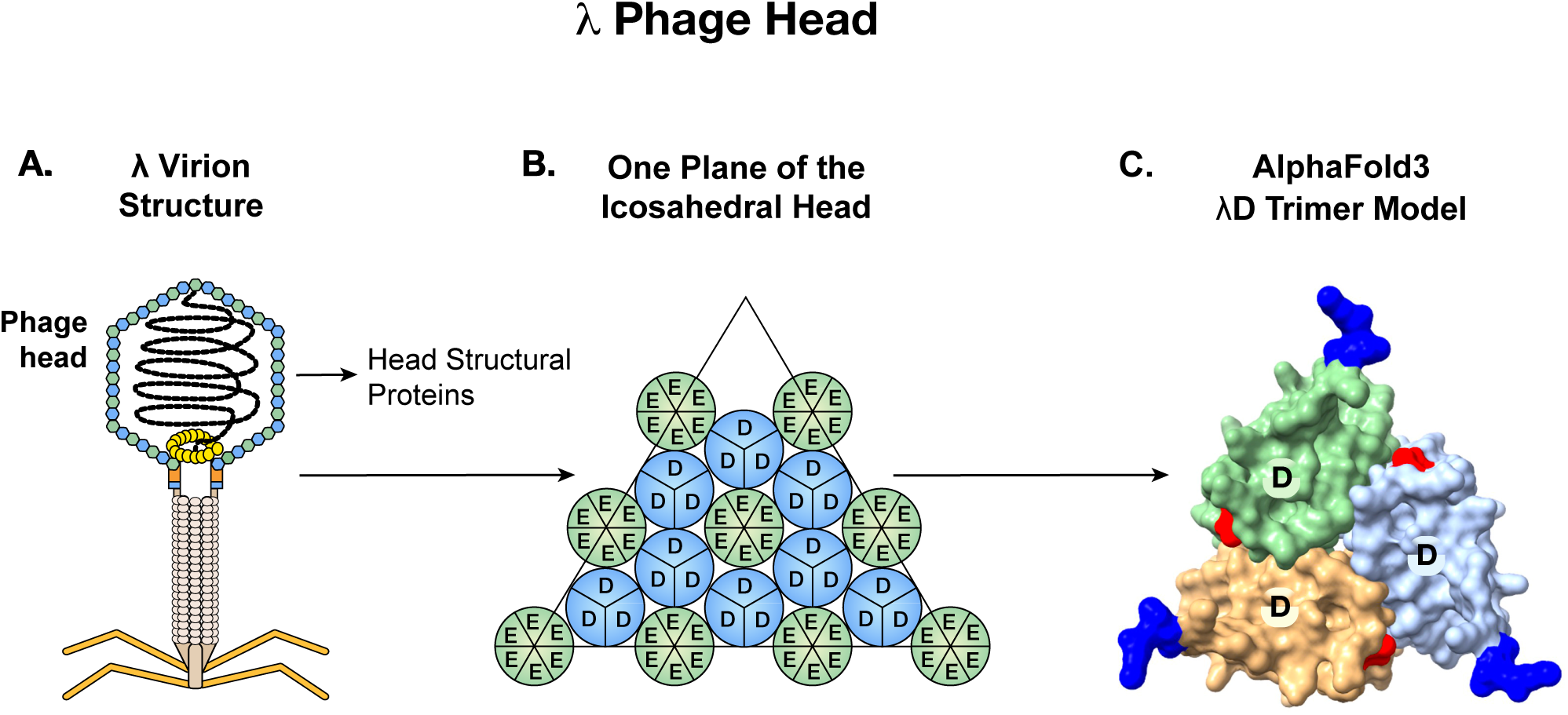
Display of antigens on bacteriophage λ. A: Graphic of phage λ structural features, highlighting the capsid proteins gpD and gpE that comprise the phage head. B: gpD trimers interspersed with gpE hexamers on the phage’s icosahedral head. Adapted from Nicastro et al., 2014. C: AlphaFold3 prediction surface model of a λD trimer with N termini in blue and C termini in red. When trimerized, both termini of λD protrude away from the head and are available to display foreign proteins.

When λ infects the bacterium *E. coli*, it may take one of two developmental paths. One is the lytic path whereupon λ usurps the host machinery to express functions required to replicate from progeny phage, lyse the cell, and release particles into the environment^4^. The alternative path is the lysogenic development following infection where the phage DNA integrates into the bacterial chromosome and produces the phage CI repressor protein to prevent expression of lytic genes. This dual-lifecycle feature allows λ to be genetically manipulated with ease while integrated in the bacterial chromosome and then induced to lytic phase to produce mature engineered phage particles.

The λ capsid is composed of two major structural proteins, gpE and gpD (gene product), each present in ∼420 copies per particle^3,5^. In the λ genome, the *D* and *E* genes are adjacent and expressed coordinately, with the gpD (gene product D) protein trimers stabilizing the primary gpE hexameric capsid structure^6^ (Fig. 1B). Functional λ particles have been made in which non-phage proteins have been fused to either the N or C terminus of gpD^1,7^. Both termini protrude away from the head, allowing the display of fusions on the surface of the capsid^8^, as seen in the AlphaFold3 generated λD trimer structure^9,10^ (Fig 1C). In most previous examples, the gpD-fusion protein was generated by expressing the D-fusion from a plasmid during infection by a λ mutant containing an inactive *D* gene. The D-fusion protein made *in trans* then assembles with the other phage components^11,12^. Here we created fusions to either terminus of the *D* gene *in vivo* on a λ prophage where the *D* fusion gene is in its normal position adjacent to gene *E* in the phage genome and expressed coordinately with gene *E* (Fig. 2).

**Fig 2.**
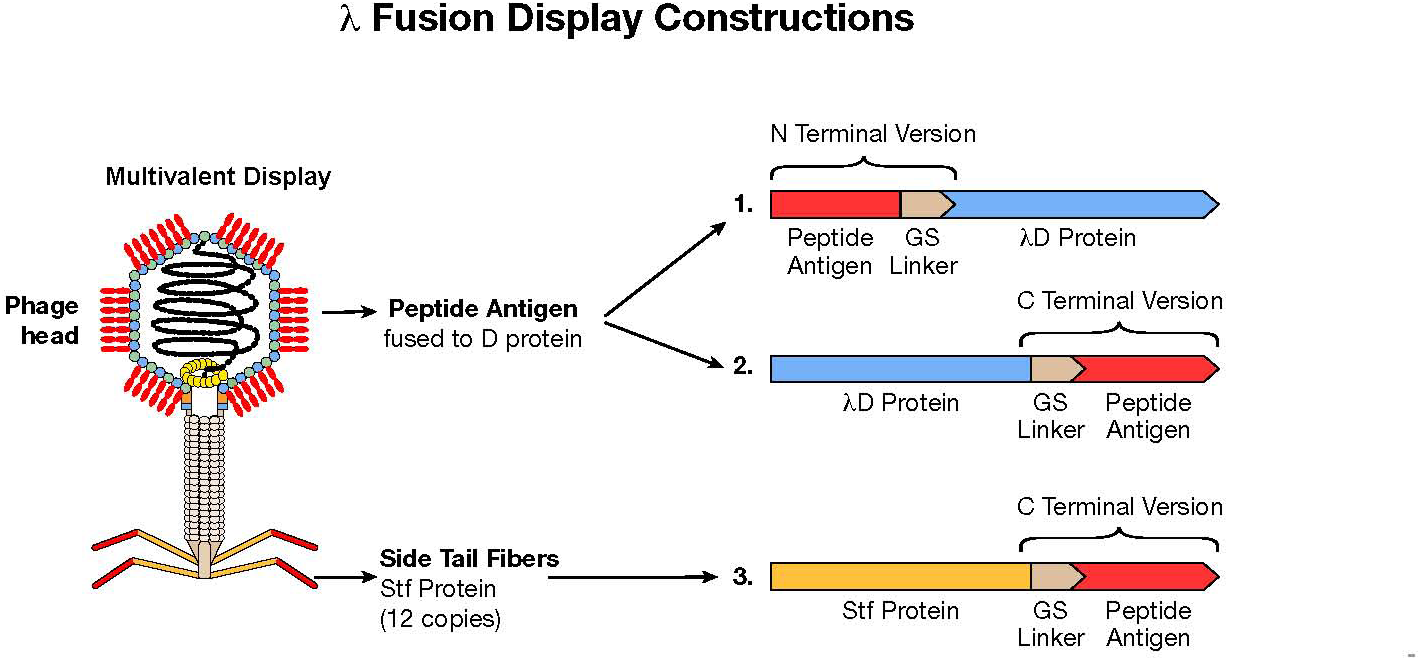
λ fusion display constructions. Graphic of phage λ with antigens fused multivalently to the λD capsid protein. Antigens can be fused either in the N– or C-terminal construction through genetic addition of a flexible glycine-serine linker and the antigen. Proteins may also be fused to the C-terminal of the Stf λ phage protein. In our construction, the Stf protein is truncated, ending at aa 386 where Stf then joins with the linker and passenger protein.

Stf of λ that attaches to the end of the phage tail can also be used to display foreign proteins, especially in longer segments than are tolerated by λD. Stf (copy number = 12) is integral for λ binding to the *E. coli* LamB receptor during infection^13^, however, the Stf protein is non-essential for λ assembly from the prophage^14^, which is the extent of our use here. Thus, we can display foreign proteins on the distal exposed carboxyl end of Stf without interrupting essential processes. Our cloning results indicate that adding protein segments at the Stf C terminus generates high-titer phage lysates for display applications.

The ability of λ to display foreign proteins has been utilized in a variety of ways: to create large peptide libraries, to identify antigenic proteins, to investigate ligand-receptor associations, to characterize proteins, to isolate peptide molecules, and to study peptide interactions, along with many other uses ^11,15–19^. The λ platform developed and tested here is an efficient display system for any of these applications. This study provides evidence for this novel display technology to be used specifically as an antigen selection system and as a vector for immune stimulation against specific passenger antigens. While other groups have reported the use of λD display to stimulate antibodies^20–24^, our modifications improve the engineering and function of the λ display vector by fusing a foreign peptide to a λ protein in its prophage state.

Several mutations have been made to facilitate the ease and quality of fusion phage engineering, prophage induction, and testing in animal studies, which are discussed below:

1. The classical λ *cI857* temperature sensitive mutation in the *cI* repressor gene causes the CI repressor to become inactive at an elevated temperature of 42° C^25^. Employing this mutation, a λ *cI857* lysogenic culture growing at 32° C can be induced to produce phage particles and lyse the cell by simply shifting the culture temperature to 42° C. This avoids using methods of induction by agents such as mitomycin or UV, which damage DNA.
2. The *Sam7* mutation was introduced to the prophage to increase phage yield. Normally the time from λ prophage temperature induction to subsequent lysis and phage release is ∼50 minutes, at which time ∼100 particles of progeny phage per cell are released. Lysis is dependent upon the phage *S* gene encoded holin, which generates micron-scale holes in the cytoplasmic membrane after ∼50 minutes of prophage induction^26^. These holes allow the endolysin to escape from the cytoplasm and degrade the cell’s peptidoglycan layer, causing complete cell lysis. If the holin protein is inactivated, then cytoplasmic endolysin cannot escape to cause lysis. However, λ continues to replicate within the cell and ultimately produces ∼1000 phage particles per cell. Here we have used the *Sam7* mutation in the prophage to prevent holin synthesis^27^. Phage lysis is caused later by the addition of chloroform or freezing and thawing the culture, thereby releasing the increased phage burst size (∼10-fold increase).
3. We deleted the 4,285-base-pair *b2* region to increase the λ phage packaging capacity. A λ virion consists of a tail and a 48.5 kb dsDNA genome tightly packaged within an icosahedral protein shell. λ phage DNA lengths that fall outside the optimal packaging range of 38-52 kilobase pairs (kb) are generally not incorporated into functional, infectious phage particles^28,29^. The *b2* region is a large, non-essential segment of the genome (approximately 5.7 kb). Deleting it does not prevent the phage from undergoing the lytic cycle. Further, its deletion increases the capacity of the vector, allowing the cloning of larger DNA fragments into the phage genome while keeping the total size within the optimal packaging range of 38-52 kb^30^.
4. The *LamB* λ receptor protein encoding gene was deleted in the host chromosome to prevent λ binding and reduced yield after lysis. Initial lysates after chloroform treatment contain bacterial cell debris including isolated external cell surface λ receptors. Receptors present in the cellular debris of the lysate can bind to λ, causing ejection of the genome into the lysate and thereby reducing the number of active phage^31^. We eliminated the latter event of yield reduction by deleting the *lamB* gene from the host *E. coli*.
5. For half-life extension of the λ phage in the mammalian circulatory system, the major capsid *E* gene has been modified from Glu158 (GAG) to Lys158 (AAG), a single amino acid change. This change enables λ particles to have a longer presence in the mammalian circulatory system, which has the potential to increase immunogenicity^32^.

A λ phage-based vaccine is an attractive therapeutic option because it confers several advantages over traditional immune-stimulating platforms. It is inexpensive to develop, and displays can be quickly engineered for countering new or evolving diseases. The phage particles themselves are stable at ambient temperatures, providing convenient shipping and storage of a λ phage display therapeutic^33^. The λ phage itself acts as an adjuvant, stimulating the body’s immune response against the high copy-number displayed antigen without requiring an added immune stimulant^22,23,34,35^. With the establishment of a reliable and effective λ platform, a variety of antigen-displaying therapies could be quickly utilized to treat or prevent disease. In this article, we describe the development of our specialized λ phage display vector and the methodology of prophage engineering and induction, as well as show data supporting the use of λ display as an antigen selection system and therapeutic vector platform.

## Results

After generating *E. coli* host strains with prophage optimized for phage display in three different constructions, we compared the stability of phage displaying a series of melanoma neo-epitopes in all three constructs. Additionally, we demonstrated the use of the C-terminal λD phage display construction for epitope library generation, antigen selection, and immune stimulation in a mouse model against the SARS-CoV-2 spike protein.

### Construct stability by titer

For the purpose of testing which of three melanoma neo-epitope display constructions would best slow the growth of melanoma tumors, we generated thirty phages displaying foreign peptides. Each of ten melanoma antigens were fused to λ in three display constructs: C-terminal Stf, N-terminal λD, and C-terminal λD. The differing titers of these three groups of phage lysates, which carry the same set of foreign peptides, illuminate phage stability by fusion construction (Table 1A-C).

**Table 1:**
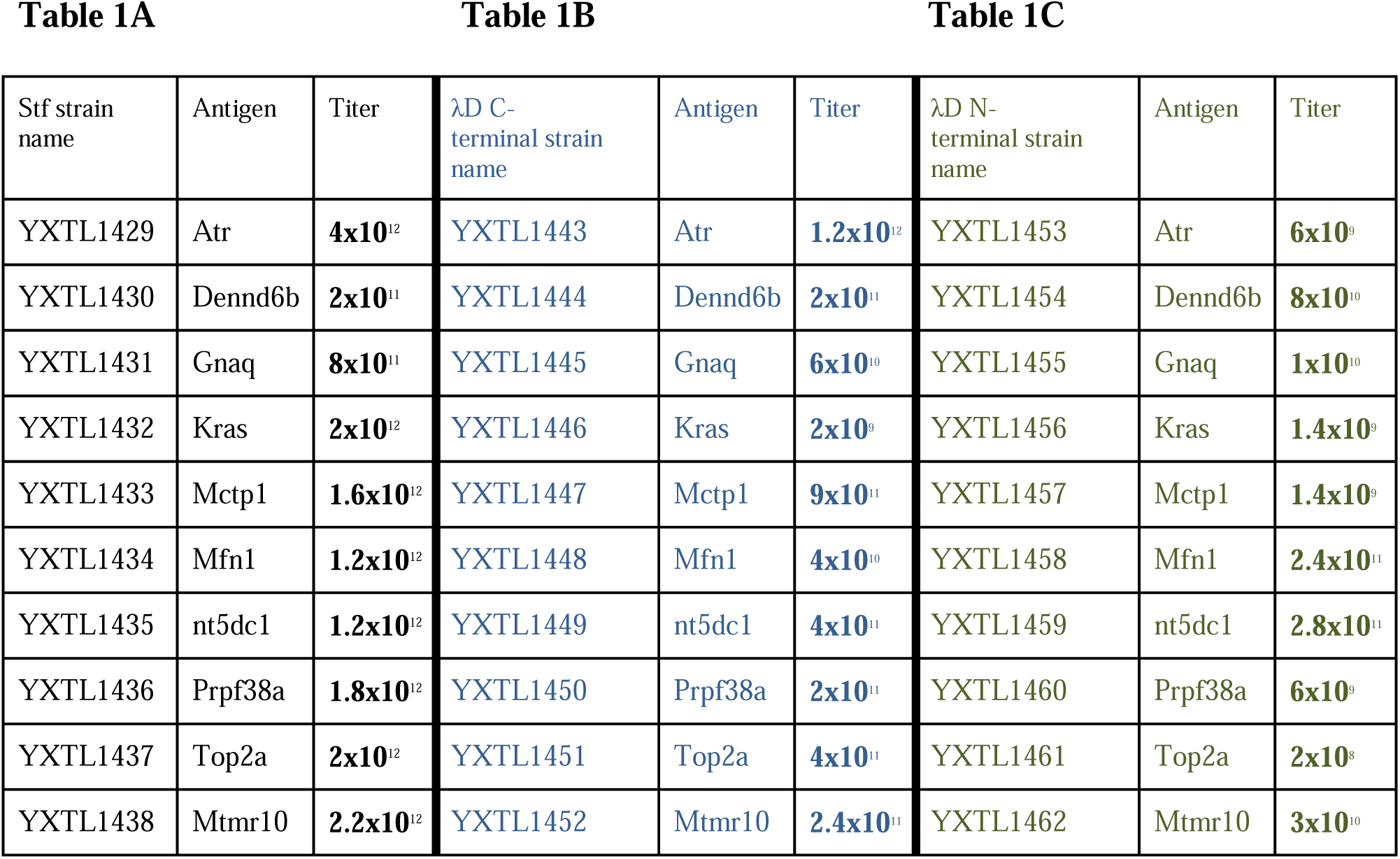
Titers of λ phage lysates displaying melanoma antigens in three engineering constructions: Stf, λD C-terminal, and λD N-terminal. Phage titers from each of thirty phage lysates prepared by induction and freeze/thaw lysis. Each phage displays one of ten antigens found in mouse melanoma cell line B2905 in three differing fusion constructions: A: Stf, B: C-terminal λD, and C: N-terminal λD. Prophage genotypes are listed in Table 2. Titers were determined by plating serial phage lysate dilutions on *supF* strain XTL1212 and counting resulting plaques after overnight incubation.

The melanoma antigens used here originated from the detection of ten distinct mutations identified in mouse melanoma cell line B2905 by whole exome sequencing^36^. To create fusion antigens, each mutation was flanked on either side by 12 amino acids from its local genetic region in cell line B2905. These ten 25-aa sequences containing melanoma mutations were recombineered into the λ prophage in the three different constructions by counter-selection recombineering described in the methods (Table 2). The fusion phage were then induced, filtered, and the lysates titered on strain XTL1212. All 30 lysates were produced with identical technique. Titering phage samples involved the serial dilution of a phage lysate, plating on a suitable bacterial host (XTL1212), and plaque counting after overnight incubation at 39° C.

**Table 2:**
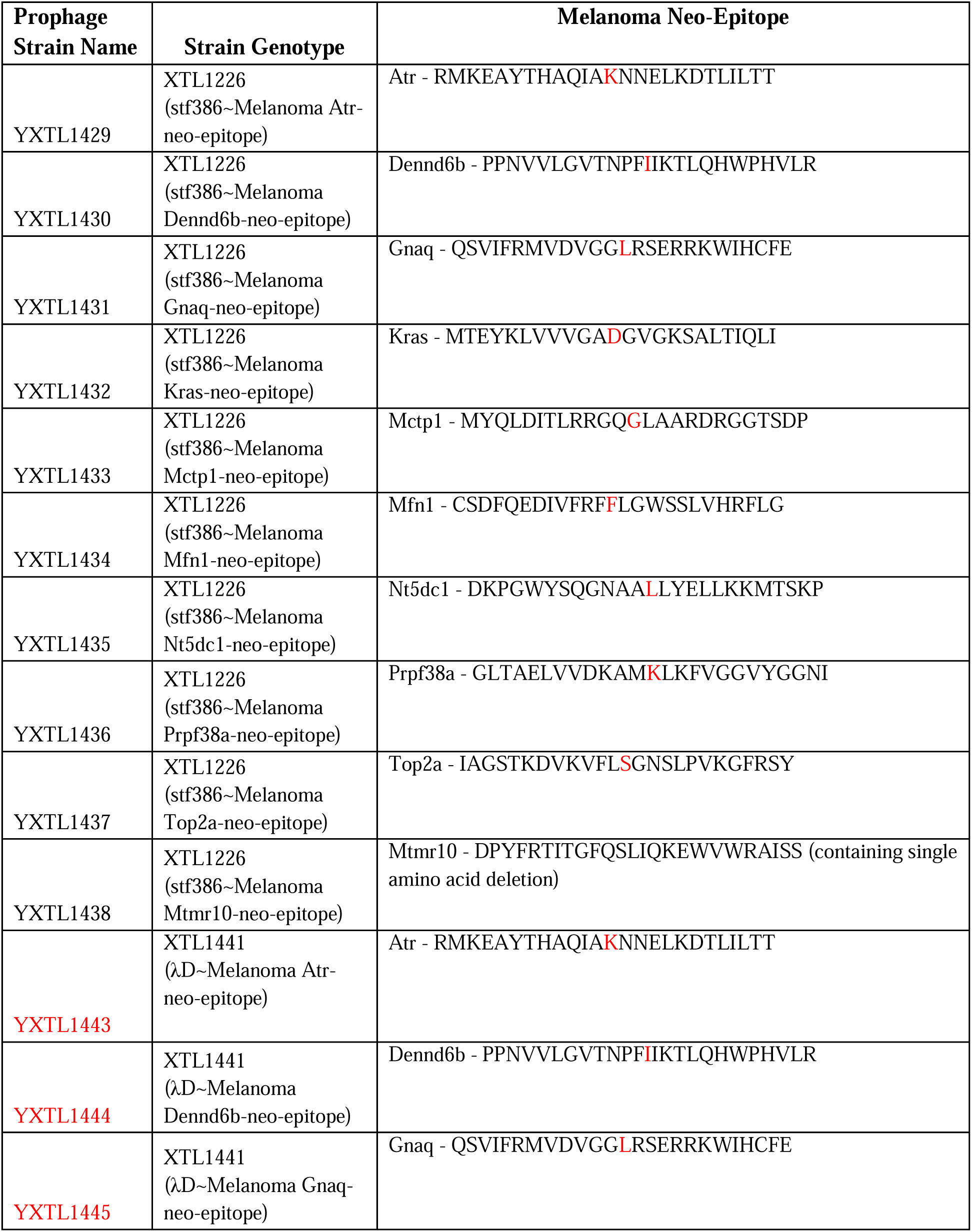
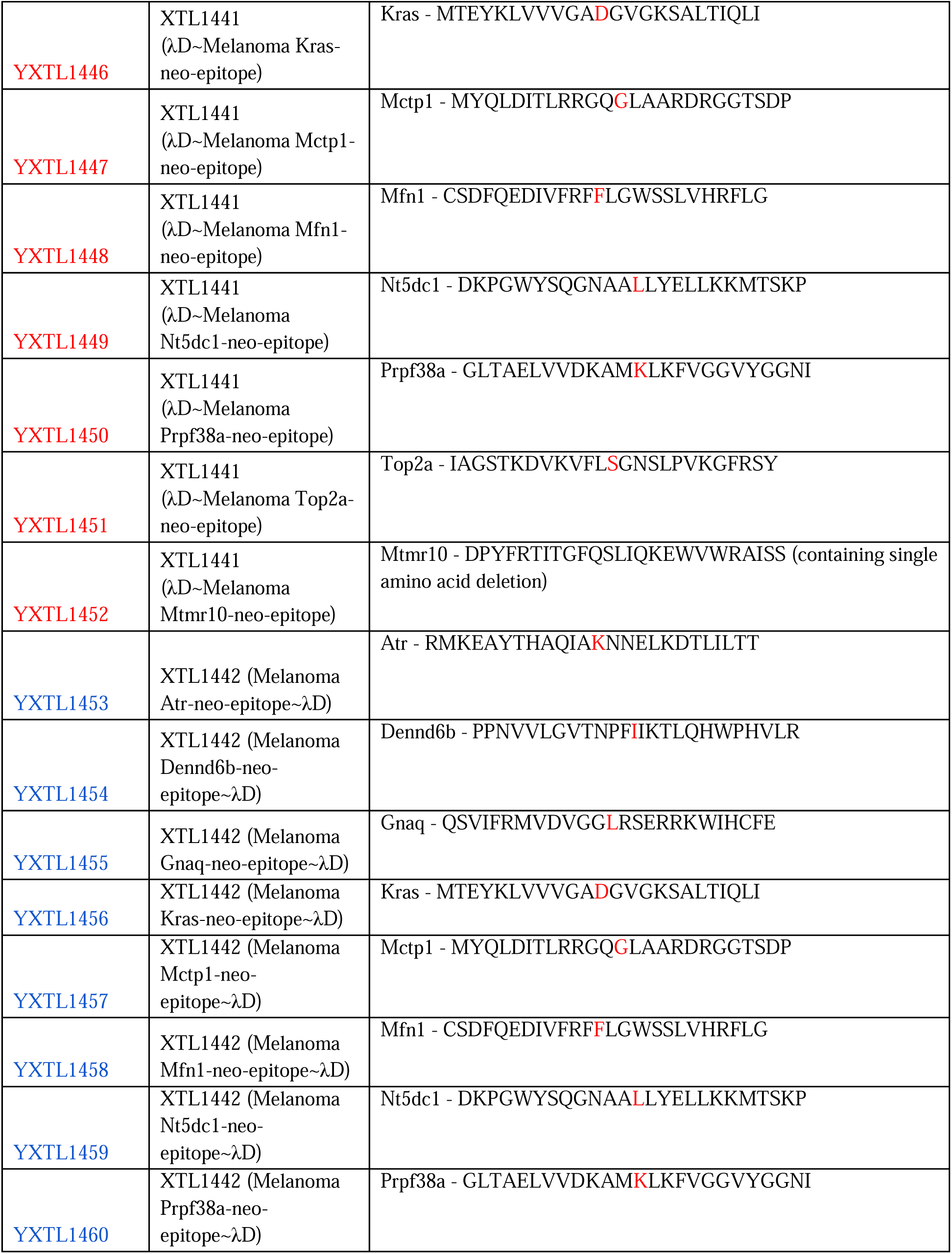
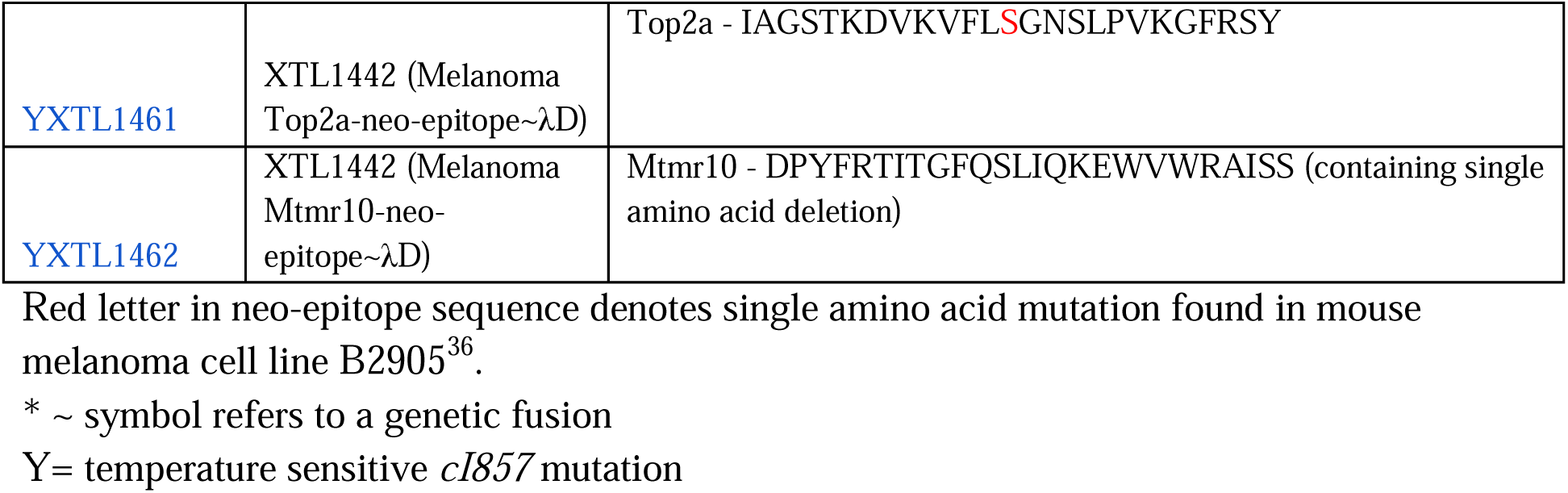
Melanoma prophage strains. Prophage strains developed to generate phage displaying each of ten melanoma neo-epitopes at three fusion locations: C terminus of Stf, C terminus of λD, and the N terminus of λD. Parental strains and melanoma epitopes listed for each prophage developed.

Stf melanoma fusions produced highest titer phages with all but two samples reaching more than 1×10^12^ pfu/ml. C-terminal λD constructions had the second highest stability with most samples titering between 10^10^ and 10^11^ pfu/ml. N-terminal lysates had the lowest average titers among the three constructions with most samples achieving 10^9^ and 10^10^pfu/ml. The N-terminal group also yielded the lowest titer among the 30 fusion phage produced with the Top2A epitope phage lysate titering at only 2×10^8^ pfu/ml. While titer among constructions may vary with the identity and length of the fusion antigen, these trends can be considered when choosing a construct for other diseases and display applications.

### *λ*D antigen detection

Here we describe the utility of the λ system using disease target COVID-19. To demonstrate the use of phage λ as an antigen selection tool, we chose to define an antigenic region of the SARS-CoV-2 spike protein through generating spike protein epitope λ displays. SARS-CoV-2, the coronavirus that causes the disease COVID-19 (Coronavirus disease 2019), has a dual subunit spike protein of 1,273 amino acids^37^. This surface glycoprotein functions to make contact with the host cell surface receptor ACE2 (Angiotensin-converting enzyme 2) and complete viral fusion to invade the cell^38^. Subunit 1 is primarily responsible for binding the human ACE2 cell-surface receptor through the spike protein’s receptor binding domain (RBD), while subunit 2 encodes proteins to accomplish the viral fusion process by which the virus gains access to the human cell^39,40^. The spike protein is the primary protein to interact with human cells, and therefore, is an ideal target for identifying a useful antigen against COVID-19.

Our goal was to define an antigenic region of the spike protein small enough to create a stable, high-titer λ∼spike protein display to use in a mouse immunogenicity study. The process by which we defined a suitable antigen was to create a series of overlapping spike protein epitopes up to 133 amino acids in length, fuse the peptides to λ through C-terminal λD display, and screen the display phages with human serum post SARS-CoV-2 infection. By western blotting, we were able to define a region of the spike protein with affinity for antibodies raised by natural SARS-CoV-2 infection. Within this larger antigenic region, we also defined a small epitope of strongest antigenicity and tested a phage displaying it in a mouse antibody-generation study.

To use λ as a detection system for a SARS-CoV-2 spike protein epitope, we first segmented the spike protein gene into 12 overlapping epitopes of ∼133 amino acids per segment and a few smaller epitopes (Table 3 and Fig. 3A). Using the methodology of λ prophage recombineering, we fused these genetic epitopes to λD in the C-terminal construction and raised a lysate of each distinct fusion phage. This created a λ display library of the spike protein, which was confirmed through blotting these phage samples and the vector phage against an anti-λD antibody (Fig. 3B). Wild-type λD has a molecular weight of 11.4^8^ kDa, seen in lane 1 (1026) in both Figure 3B blots. Fusions to λD increase the overall size of the protein, raising the location of the band on the western blot to reflect the higher kDa weight of the fusion protein. Therefore, λD carrying fusions of various sizes appears as a higher band than wild-type λD in all lanes besides the vector, which contains unmodified λD protein only.

**Fig 3.**
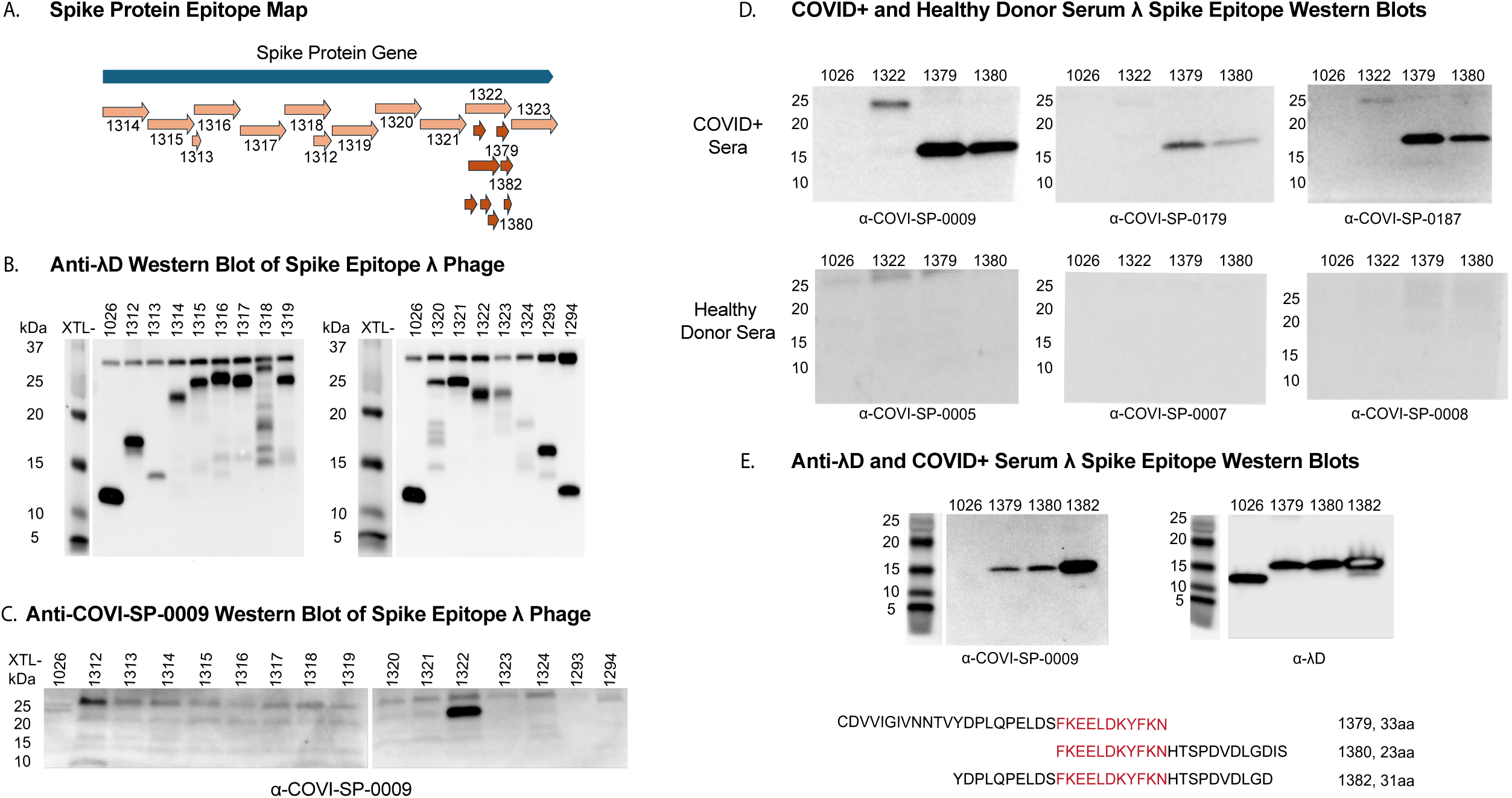
Spike protein antigen detection by human SARS-CoV-2 post-infection serum. A: Genetic map of the SARS-CoV-2 *spike* gene, divided into major epitopes for fusion to phage for antigen detection by human serum. Arrows beneath epitope XTL1322 show the segmentation of XTL1322 into 8 smaller, overlapping epitopes. B: Western blot showing a panel of 15 spike protein epitope fusion phages and a vector phage blotted against an anti-λD antibody. Bands reflect λD protein at different molecular weights depending on the size of the fusion protein. C: Western blot of the same 15 spike protein epitope fusion phage and vector phage blotted against human serum after SARS-CoV-2 infection. Band reflects serum antibody binding to phage-displayed antigen. D: Western blots where two spike protein small epitopes displayed on λ are recognized by COVID+ serum antibodies and unreactive to serum from healthy donors. Samples: vector λ phage (lane 1), large epitope XTL1322 (lane 2), and small epitopes XTL1379 (lane 3) and XTL1380 (lane 4). E: Western blots of vector λ phage (lane 1) and small epitopes XTL1379 (lane 2), XTL1380 (lane 3), and XTL1382 (lane 4). These four samples were blotted against the SARS-CoV-2-infected serum (left) and an anti-λD antibody (right). Three smaller overlapping epitopes were found to be antigenic against COVID+ serum: XTL1379, XTL1380, and XTL1382. Their sequences are reported, aligned to show overlapping amino acids. A molecular marker is shown to the left of each blot.

**Table 3:**
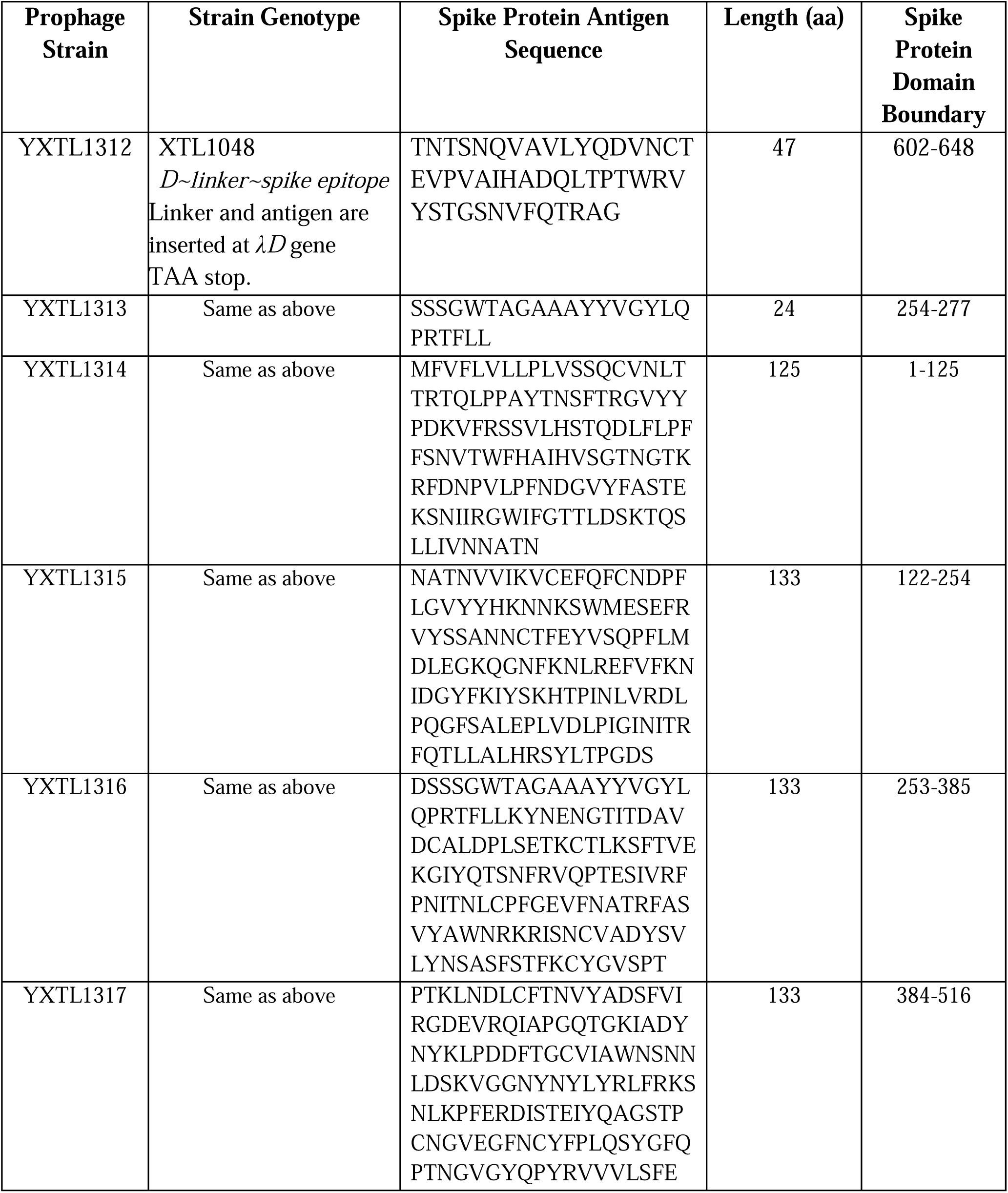
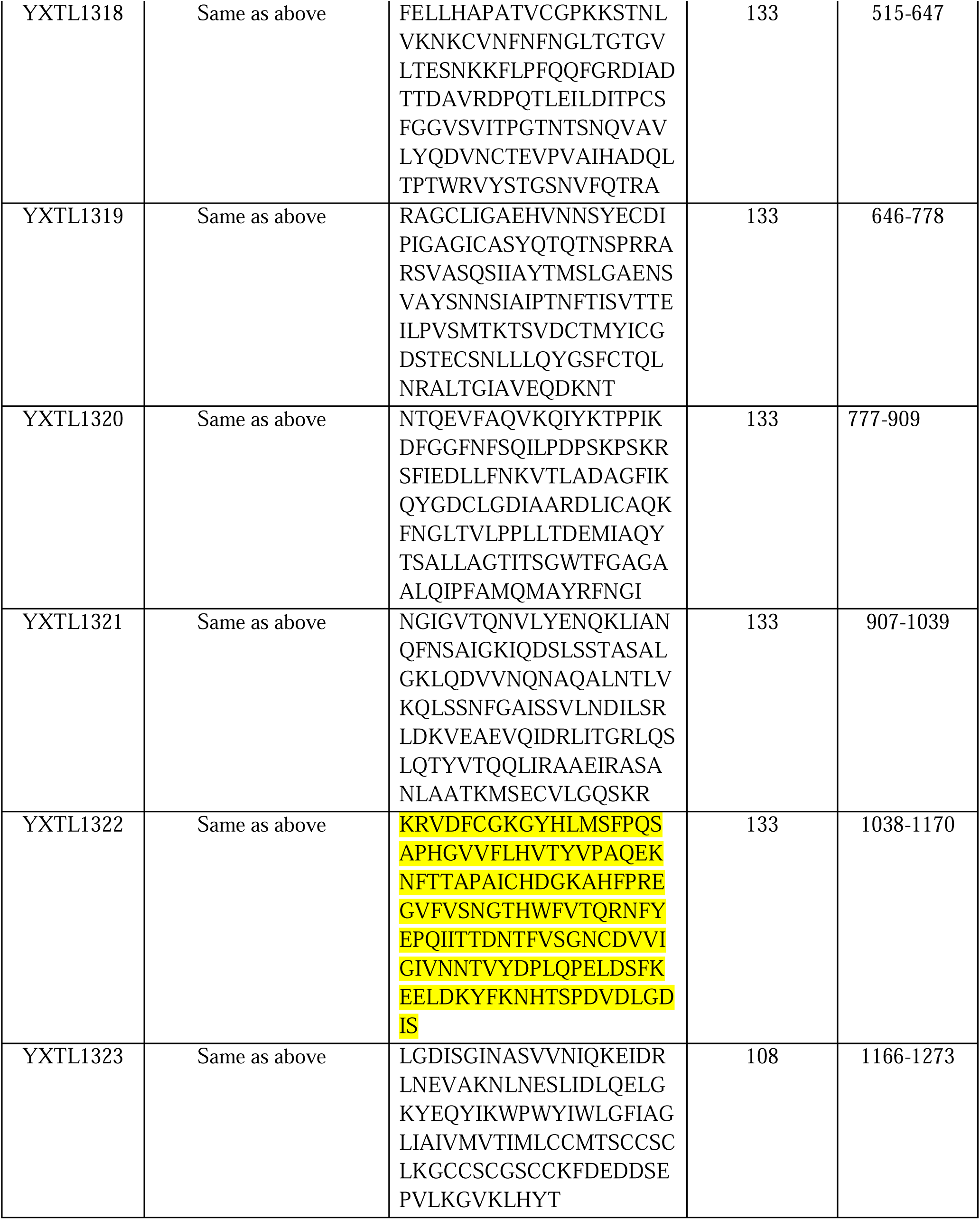
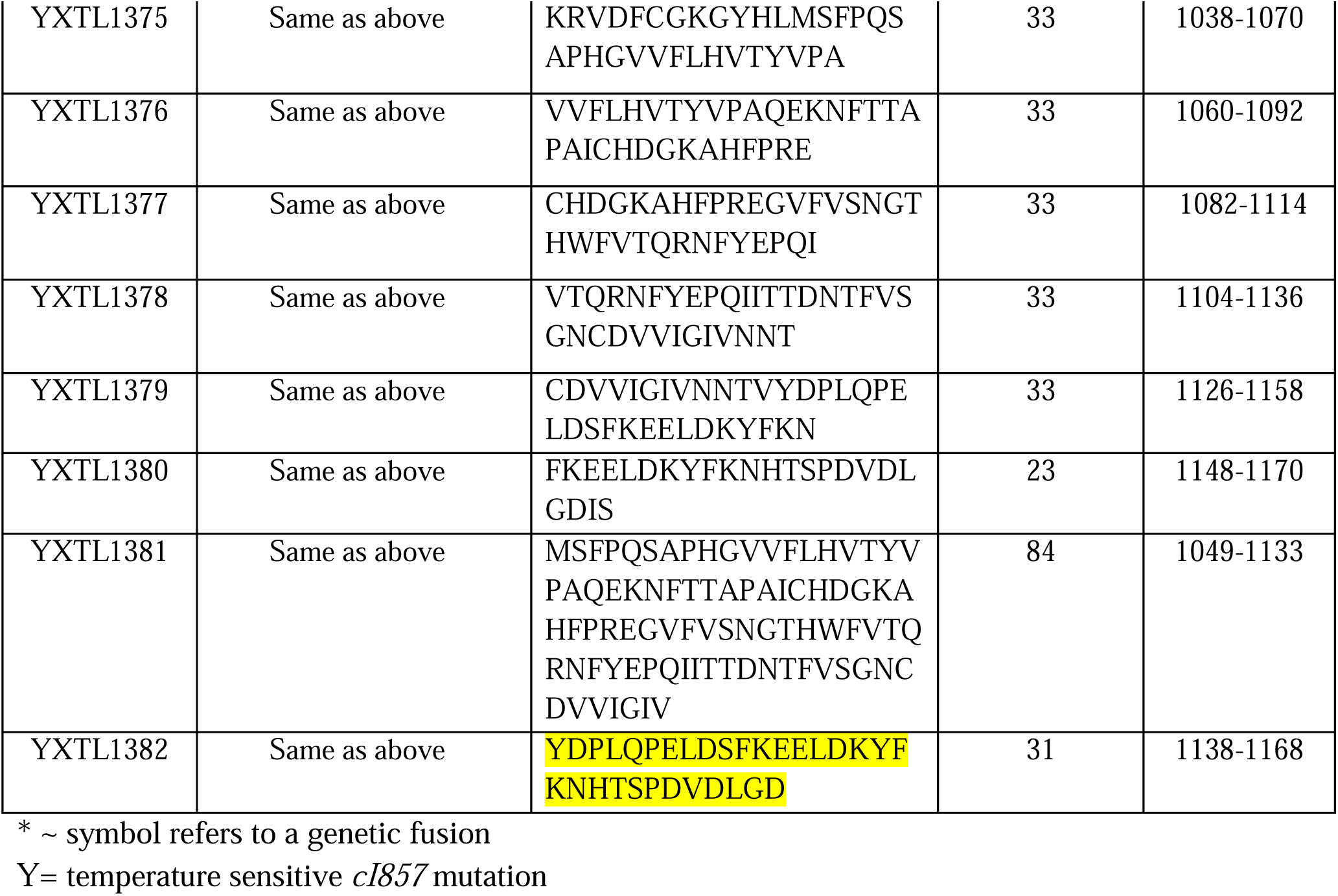
SARS-CoV-2 spike protein prophage strains and their displayed epitopes. Prophage strains developed for use in detecting a suitable SARS-CoV-2 spike protein antigen. Each strain is a modification of YXTL1048, containing a partial sequence of the spike protein fused to the C-terminal of the prophage’s λ*D* gene with a single unit linker in the sequence: GGGGS. Highlighted sequences are those displayed antigens which produced a strong response against COVID+ donor serum.

Also included in Figure 3B are phages displaying an epitope from the SARS-CoV-2 E protein (XTL1324) and two epitopes from the SARS-CoV-2 N protein (XTL1293 and XTL1294). These epitope candidates were not useful antigens and were not investigated further. The rest of the antigen investigation narrowed to only the spike protein epitopes.

This library of spike protein phage-displayed epitopes was then blotted against human serum collected after a SARS-CoV-2 infection (Fig. 3C). Only phage strain XTL1322, displaying a 133 aa peptide from the second subunit of the spike protein, reacted with COVID+ serum. This epitope consists of spike protein amino acids 1,038-1,170, which includes the subunit 2 connector domain (CD), open reading frame 1069-1162, and the first 7 amino acids of heptad repeat 2 (HR2). Both the CD and HR2 regions assist in viral fusion to the mammalian cell membrane after the spike protein receptor binding domain contacts the ACE2 cell surface receptor^37^.

Fusion protein XTL1322 created a low titer lysate, likely due to decreased stability because of the large size of the 133 aa fusion protein. To construct a more stable phage, epitope XTL1322 was further dissected into overlapping DNA sequences encoding smaller peptide fragments (Fig. 3A). These DNA fragments were recombined in-frame at the 3’ end of the λ*D* gene as described previously. While many smaller XTL1322 epitopes were tested against human COVID+ serum, only XTL1379 and XTL1380 reacted. These two epitopes were further probed against serum samples from three COVID+ patients and three healthy donors (Fig. 3D). All COVID+ sera reacted with the phage-displayed epitopes, while the healthy sera did not react with any of the phage samples.

The peptides displayed by these two phage strains overlap by 11 amino acids (Fig. 3E). Upon sequence alignment with spike proteins of different variants such as alpha, beta, delta, omicron, as well as SARS-CoV-1, we found that this 11 amino acid overlapping segment between XTL1379 and XTL1380 was conserved among all other variants of SARS-CoV. We proceeded to reengineer a new hybrid strain XTL1382 that displayed the overlapping 11 amino acid peptide flanked by the consecutive 10 amino acids at either end of the peptide. When equal plaque forming units of λ displaying XTL1382, 1380, and 1379 were again probed with COVID+ serum (COVI-SP-0009) (Fig. 3E), we found that the newly engineered hybrid strain XTL1382 strongly reacted with the antibody present in the serum as indicated by the band intensity in Fig. 3E, left. Fig. 3E, right, shows the same phage samples blotted against an anti-λD antibody, demonstrating correct fusion identity of display phages compared to the vector phage.

While any of the three small displayed epitopes (XTL1382, XTL1380, and XTL1379) were strong candidates for use in a mouse immunogenicity study, we chose to test phage displaying XTL1382 since it gave the most robust signal against COVID+ serum antibodies.

### Mouse immunogenicity study with *λ*D display phage

XTL1382 λD epitope fusion phage display lysate was tested in a mouse study for the phage’s ability to stimulate antibodies against the displayed spike epitope. Groups of mice were immunized with fusion phage and vector phage three times over the course of the 49-day study (Fig. 4A). Post-injection serum samples were tested for antigen-specific antibody production through both ELISA and western blotting.

**Fig. 4.**
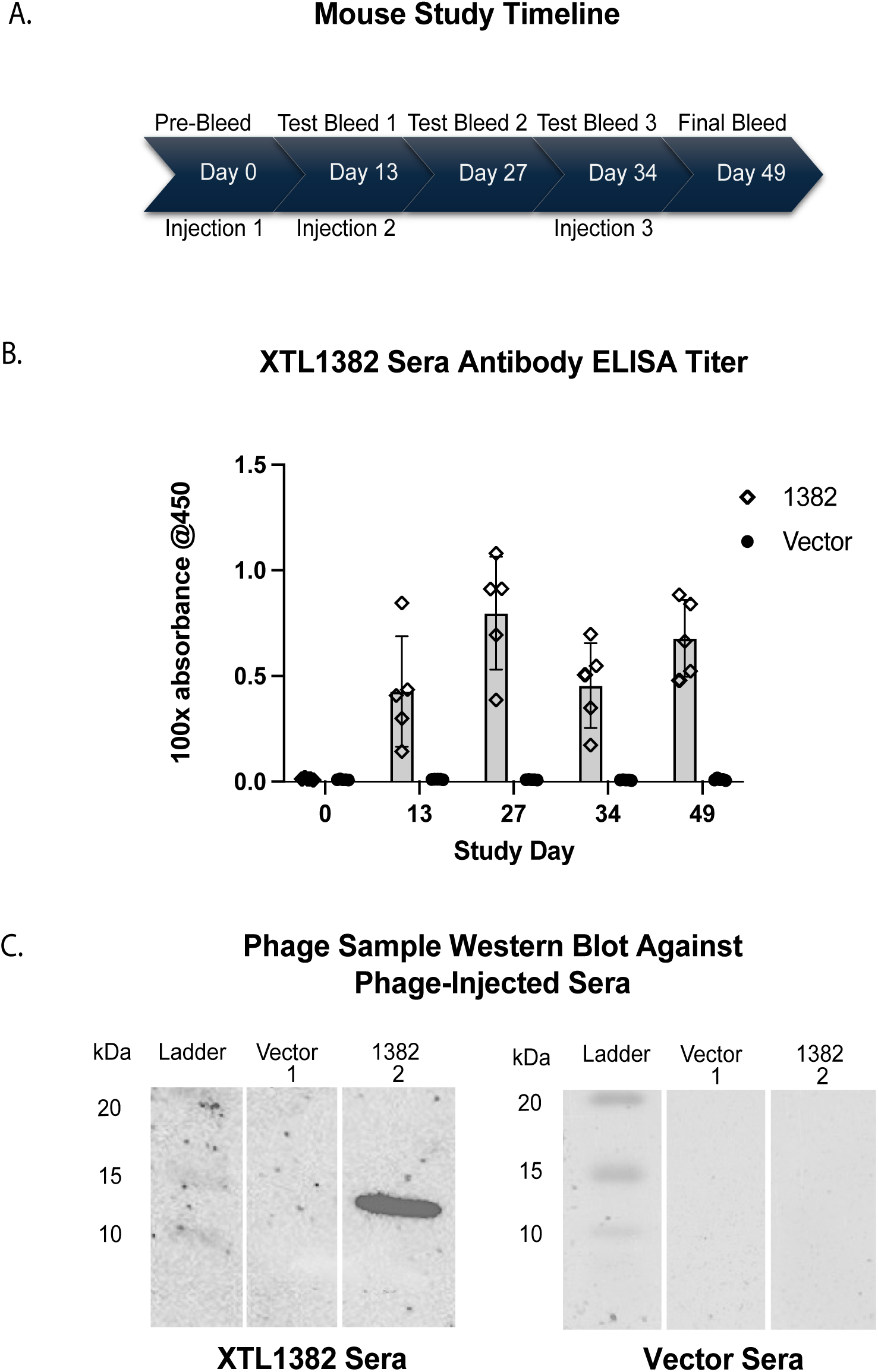
Antibody generation from XTL1382 displayed on λ phage. A: Timeline of mouse immunogenicity study. Vector and XTL1382 fusion phage were injected in groups of five mice through an IP route on Days 0, 13, and 34. Sera was gathered on Days 0, 13, 27, 34, and a final bleed on Day 49. B: ELISA antibody titer absorbance data at 100x dilution, testing vector and fusion phage sera groups (n=5) against the SARS-CoV-2 synthetic XTL1382 peptide. Antibody titer was measured at each time point over the course of the study with three readings averaged per mouse and five mice per bar. C: Western blot of vector (lane 1) and XTL1382 phage (lane 2) blotted against Day 49 XTL1382-injection serum (left) and Day 49 vector-injection serum (right).

Sera from mice injected with XTL1382 fusion phage demonstrated high XTL1382-specific antibody titers at 100x dilution two weeks after initial vaccination (Fig. 4B). Titer increased further after Day 13 injection but dipped between the second and third injection on Day 34. However, upon re-injection on Day 34, titers rose again to a high level on the final bleed day. Pre-bleed antibody titers as well as all vector phage-injected sera samples remained at a background absorbance level, indicating the specificity of antibody response detected in the experimental serum group.

Western blots confirmed that phage displaying the XTL1382 antigen created XTL1382-specific antibodies in mice (Fig. 4C). When vector (lane 1) and XTL1382 phage (lane 2) were blotted with serum from a mouse that received the XTL1382 injection (left), the XTL1382 phage lane showed a strong band at the expected λD fusion protein region of ∼15 kDa. The vector phage lane was blank, indicating no protein binding with mouse serum antibodies. As a control, both phage samples were blotted against serum from a mouse injected with vector phage (right). No bands appeared in either phage lane, indicating a lack of antibody presence specific to either the vector or XTL1382 fusion phage λD protein.

This study was a confirmation of a previous mouse study that likewise injected XTL1382 display phage into a group of mice (N=3). That study found similar antibody levels in response to phage display injection and yielded the same western blot results confirming presence of XTL1382-specific antibodies (Fig. S1A-C). These two antibody-detection methods, ELISA and western blotting, show XTL1382-specific antibody production in mice from an injection of λ displaying the epitope, demonstrating the use of the λ vector for immune-stimulation.

## Discussion

The production of an optimized λ phage vector for antigen detection and immune stimulation involved the evolution of strain SJ_XTL175 by several steps of genetic recombineering through the *tetA-sacB* cassette counter-selection method described in materials and methods^41^. Our system uniquely utilizes this technique to make a direct gene fusion in the prophage chromosome. The mutations to the platform maximize phage titer, increase particle stability, and expedite the engineering process. Additionally, our methodology streamlines prophage induction through the heat-inducible genetic switch and purifies the samples through tangential flow filtration. With ready-to-modify strains for any of the three fusion constructions, a stable lysate of a phage displaying a foreign protein can be made in a week’s time. This simple and rapid methodology allows for the use of λ phage display for many applications.

The display platform is suitable for foreign protein display in three constructions: the C or N terminus of the λD capsid protein or the C terminus of the Stf protein. We have shown that protein fusions to any of these locations can produce phage display lysates. When creating a library of phage displaying melanoma antigens, the Stf fusion constructions yielded the highest titer phage lysates, followed by the C-terminal λD fusion constructions (Table 1). N-terminal λD fusions yielded the lowest virion stability and titer among the three fusion construction options, which may limit its use. A possible structural explanation for this stability difference is that the first 14 N terminus residues of λD were shown through crystal structure to be disordered and positioned near the three-fold λD trimer axis^8^. Conversely, C-terminal residues had a stable confirmation and protruded from the protein away from the trimer axis. The N terminus’ high flexibility, lack of consistent conformation, and position near an essential assembly point may contribute to particle instability when fusing to this location. N-terminal fusion lower titer may be potentially mitigated by inducing a supply of helper wild-type λD from the host chromosome through arabinose addition during phage assembly (Fig. 6). Additionally, this titer stability trend may differ when fusing other peptides from those tested here. While Stf fusion constructs yielded the highest titer phage lysates, λD fusions package 35x more antigens per phage particle. This difference in protein copy number per phage particle (∼420 for λD vs. 12 for Stf) may make λD fusions, particularly in the C-terminal construction, more attractive for certain applications where a high copy number is desired.

Here, we created a displayed spike protein library to detect segments of the SARS-CoV-2 spike protein which had affinity for human COVID+ serum antibodies. Through the process of library generation and antigenicity testing, we were able to detect a 133 aa region of the spike protein, XTL1322, which consistently reacted against human COVID+ serum (Fig. 3). This technique of generating a foreign peptide display library and running antibody-protein affinity tests, also called biopanning, could be easily replicated with the λ system to identify proteins of interest for a variety of diseases. Additionally, as shown here by the selection of small epitope XTL1382 within the larger XTL1322 epitope, the λ system can also define specific regions of high antigenicity within larger proteins. Initially, we selected overlapping epitopes XTL1379 and XTL1380 for their antibody affinity observed on COVID+ serum western blots. Upon investigation of the 11aa overlapping sequence between the two and discovering that this was a highly conserved coronavirus region, we created an epitope centering this sequence with flanking arms. The resulting epitope had the strongest affinity for COVID+ serum antibodies. This method of testing overlapping protein segments could be useful for defining regions of high antigenicity in other disease targets.

To demonstrate λ’s ability to stimulate a strong and specific antibody response against a selected antigen, we performed a mouse study, injecting the same display phage used to detect spike protein antigen XTL1382 along with vector phage into groups of mice (Fig. 4). Antibody levels detected by ELISA and western blotting from study sera indicate that the λ-displayed antigen was able to stimulate a rapid, strong, and specific antibody titer against the XTL1382 peptide. Importantly, this immune-stimulating effect from λ display injections was achieved without the use of any additional adjuvant.

The COVID-19 antigenicity study described here is being further investigated by testing the mouse antibodies created by these and other spike protein antigens for their neutralizing ability against SARS-CoV-2 infection. Through several ongoing studies fusing other disease therapeutics to the three fusion locations, we are continuing to investigate the best engineering construction and mouse study conditions of the λ phage display platform, including exploring applications of Stf display, comparing N– and C-terminal λD fusion, testing linkers of differing lengths, and determining the most efficacious routes of injection.

The three uses reported here: library generation, antigen detection, and immune stimulation, are only a few of the possible applications of this λ display technology. With the ease, timeliness, and low cost of this antigen display method, the potential uses reach as far as display and vector delivery needs expand.

## Acknowledgment

This research was supported by the Intramural Research Program of the National Institutes of Health (NIH), National Cancer Institute (NCI), and the Center for Cancer Research. (CCR). The contributions of the NIH author(s) were made as part of their official duties as NIH federal employees, are in compliance with agency policy requirements, and are considered Works of the United States Government. However, the findings and conclusions presented in this paper are those of the author(s) and do not necessarily reflect the views of the NIH or the U.S. Department of Health and Human Services.

We would like to thank Hong Zhou for performing ELISA assays and collecting data found in supplemental Figure 1. We also thank all other members of our lab for their helpful assistance.

## Methods

### Recombineering

All genetic changes made to the host strain (Table 4A) and its prophage λ (Table 4B) were made by the same general method of recombineering^41^. All primers used to make these genetic changes and check accuracy are reported in Table 4C. For all recombineered strains, a *tetA-sacB* cassette was inserted at the site on the bacterial chromosome or prophage genome to be deleted or altered by selecting for tetracycline resistance and ensuring that the drug-resistant recombinants had also become sensitive to sucrose because of the *sacB* gene (Fig. 5). Using recombineering technology, the *tetA-sacB* cassette is replaced using PCR fragments, gene blocks, or single strand oligos containing the coding sequence of the antigenic genes to be fused to λ*D* or *stf*. The single strand DNA oligonucleotide is flanked by the homologous sequences to target the chromosome or prophage genome for recombination. These sucrose-resistant recombinants are tetracycline sensitive and are engineered to fuse the antigenic gene to the *D* or *stf* gene in the same reading frame. When placed at the λ*D* N terminus, the fusion DNA has an upstream 5’ ATG start codon, and when placed at the λ*D* C terminus, the fusion DNA has a downstream 3’ TAA stop codon. Additional DNA coding sequences are included to generate a flexible linker sequence between the λD or Stf protein and the foreign protein from one to three repeats of the amino acid sequence GGGGS^42^. Once fusions are generated, the λ*D*-fusion or *stf-*fusion segments are amplified by PCR and sequenced to verify the accuracy of the new gene fusions.

**Fig 5.**
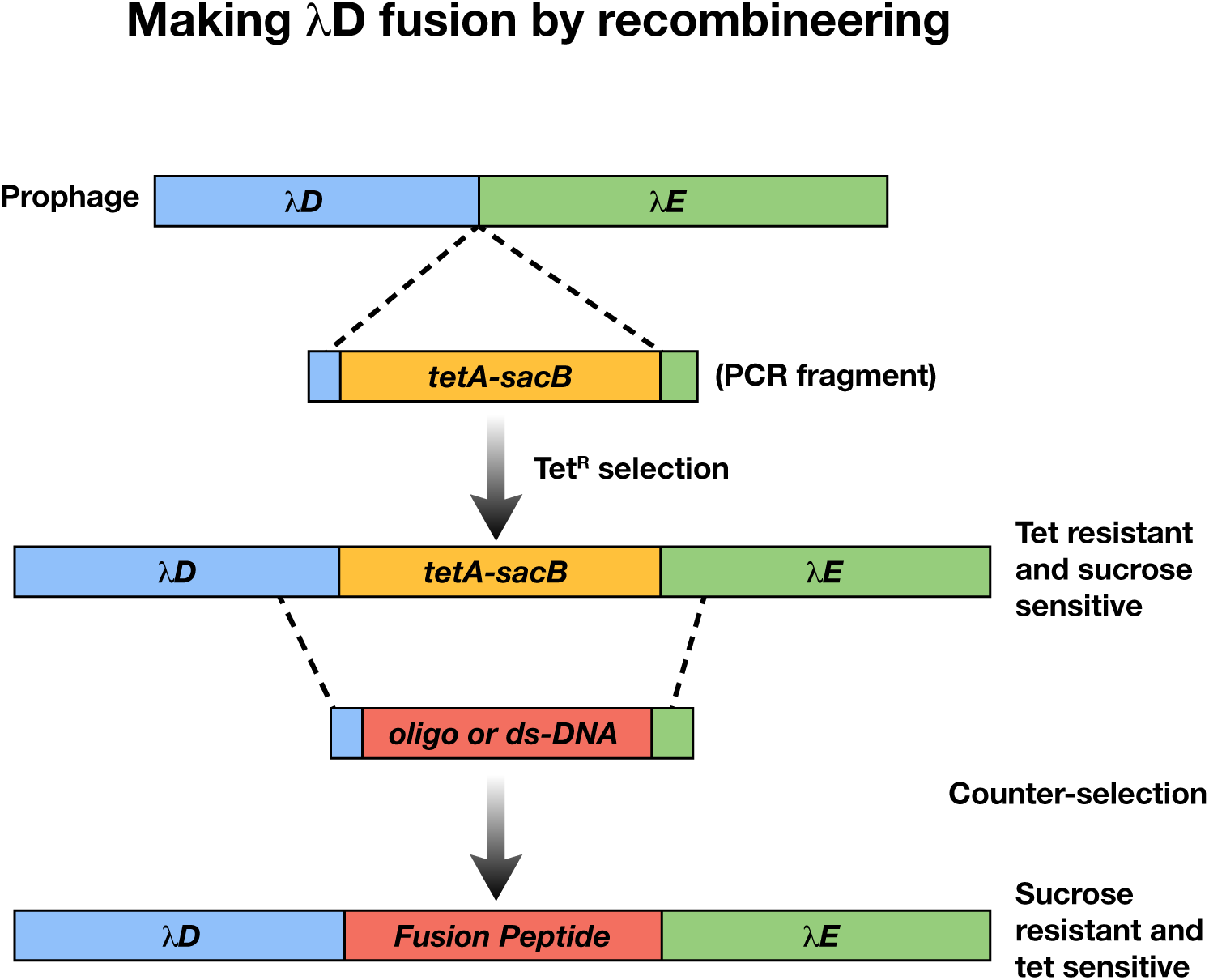
Recombineering by counter-selection. A: The λ prophage genome is modified by inserting a *tetA-sacB* cassette either at the C or N terminus of *λD*. In the figure example, the cassette is inserted at the C terminus between *λD* and *λE* genes. This modification creates tet-resistant, sucrose-sensitive colonies. A sequence of a foreign peptide can then be inserted to replace the *tetA-sacB* cassette, creating counter-selection. This resultant modified strain contains the foreign peptide and is tet sensitive and sucrose resistant. Clones can be selected by growth on sucrose plates.

**Figure 6.**
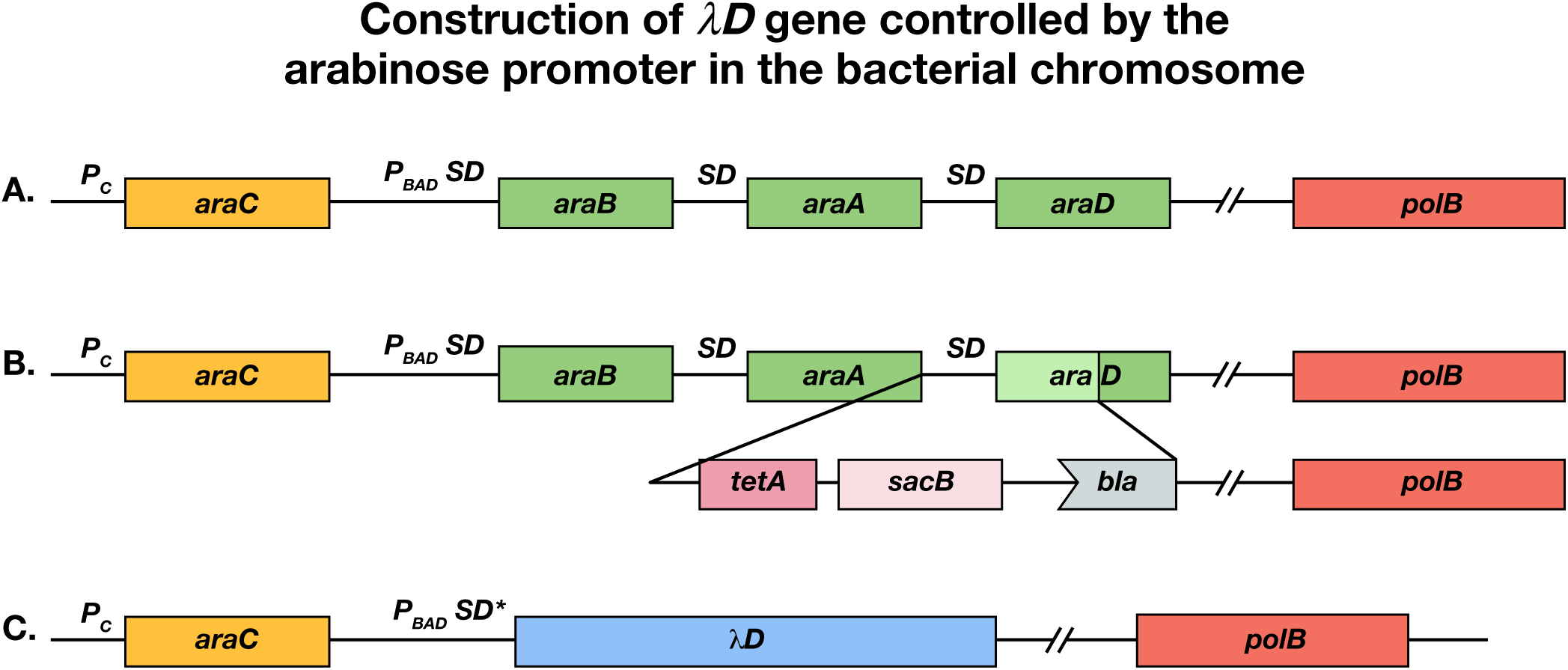
Construction of *λD* gene controlled by the arabinose promoter *pBAD*. (a) The wild-type arabinose operon *araC araBAD* with the *polB* gene of DNA Pol II downstream. SD indicates the Shine-Dalgarno ribosome binding site. (b) Insertion of *tetA-sacB* cassette downstream of *araA* to replace part of *araD*. (c) Replacement of SD *araB, araA, cat, tetA, bla* and remainder of *araD* with the *λD* gene and an optimized SD* ribosome binding site.

**Table 4A:**
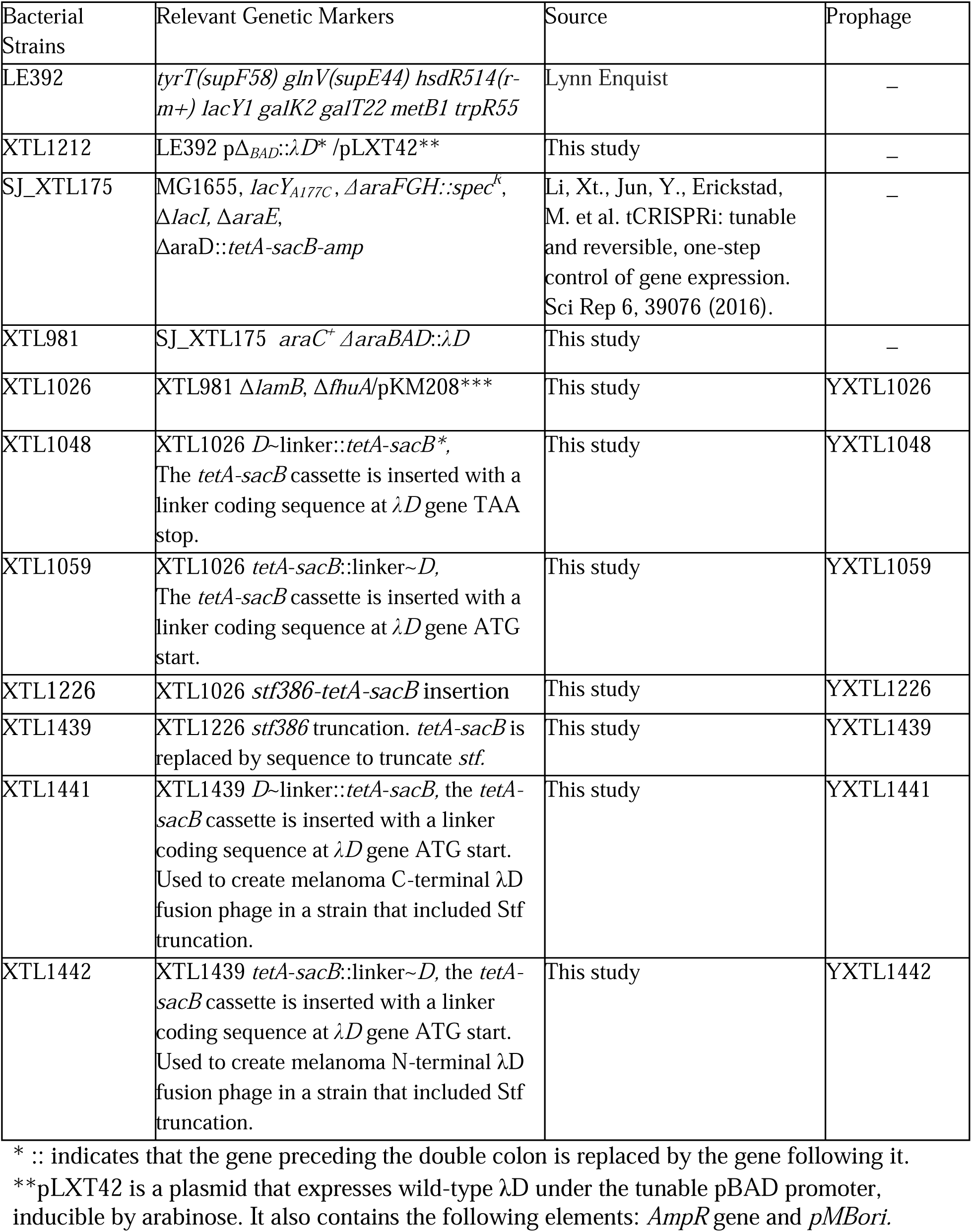
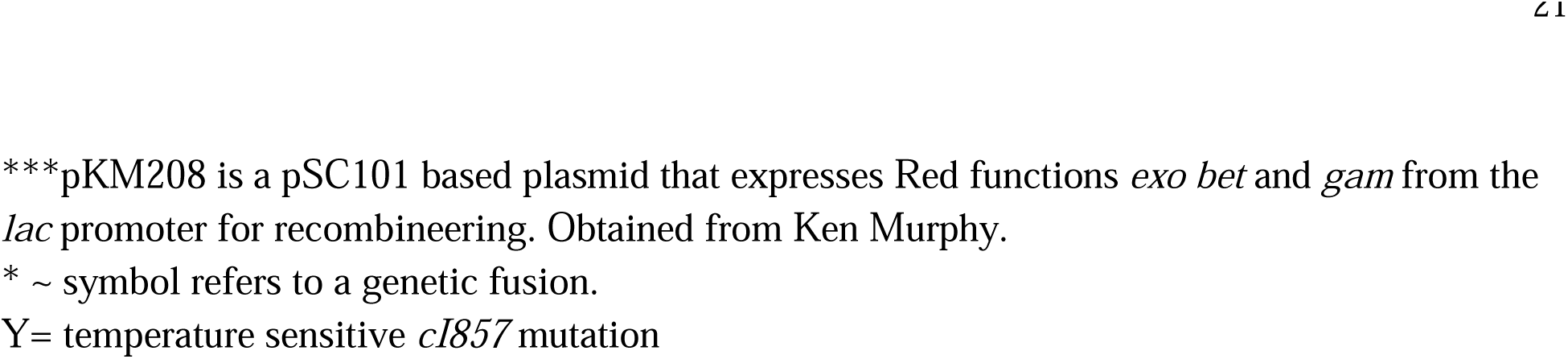
*E. coli* strains used and/or generated here. Bacterial strains generated in the evolution of a λ display platform. Table includes the initial parental strain, each new strain generated to modify the bacterial host, each strain containing modifications to λ prophage within the host, and the final strains used to create antigen-displaying phages in this study.

| Bacterial Strains | Relevant Genetic Markers | Source | Prophage |
| --- | --- | --- | --- |
| LE392 | *tyrT(supF58) glnV(supE44) hsdR514(r-m+) lacY1 galK2 galT22 metB1 trpR55* | Lynn Enquist | – |
| XTL1212 | LE392 p$\Delta_{BAD}::\lambda D^*$ /pLXT42** | This study | – |
| SJ_XTL175 | MG1655, *lacY<sub>A177C</sub>*, Δ*araFGH::spec<sup>R</sup>*, Δ*lacI*, Δ*araE*, Δ*araD::tetA-sacB-amp* | Li, Xt., Jun, Y., Erickstad, M. et al. tCRISPRi: tunable and reversible, one-step control of gene expression. Sci Rep 6, 39076 (2016). | – |
| XTL981 | SJ_XTL175 *araC<sup>+</sup>* Δ*araBAD::\lambda D* | This study | – |
| XTL1026 | XTL981 Δ*lamB*, Δ*fhuA*/pKM208*** | This study | YXTL1026 |
| XTL1048 | XTL1026 *D* ~linker:: *tetA-sacB* *,<br>The *tetA-sacB* cassette is inserted with a linker coding sequence at $\lambda D$ gene TAA stop. | This study | YXTL1048 |
| XTL1059 | XTL1026 *tetA-sacB*::linker~*D*,<br>The *tetA-sacB* cassette is inserted with a linker coding sequence at $\lambda D$ gene ATG start. | This study | YXTL1059 |
| XTL1226 | XTL1026 *stf386-tetA-sacB* insertion | This study | YXTL1226 |
| XTL1439 | XTL1226 *stf386* truncation. *tetA-sacB* is replaced by sequence to truncate *stf*. | This study | YXTL1439 |
| XTL1441 | XTL1439 *D*~linker::*tetA-sacB*, the *tetA-sacB* cassette is inserted with a linker coding sequence at $\lambda D$ gene ATG start. Used to create melanoma C-terminal $\lambda D$ fusion phage in a strain that included Stf truncation. | This study | YXTL1441 |
| XTL1442 | XTL1439 *tetA-sacB*::linker~*D*, the *tetA-sacB* cassette is inserted with a linker coding sequence at $\lambda D$ gene ATG start. Used to create melanoma N-terminal $\lambda D$ fusion phage in a strain that included Stf truncation. | This study | YXTL1442 |

\* :: indicates that the gene preceding the double colon is replaced by the gene following it.
\*\*pLXT42 is a plasmid that expresses wild-type $\lambda D$ under the tunable pBAD promoter, inducible by arabinose. It also contains the following elements: *AmpR* gene and *pMBori*.
\*\*\*pKM208 is a pSC101 based plasmid that expresses Red functions *exo bet* and *gam* from the *lac* promoter for recombineering. Obtained from Ken Murphy.
\* ~ symbol refers to a genetic fusion.

**Table 4B:** λ prophage used and/or generated here. Complementary prophage strains to Table 4A, listing the genetic modifications of each prophage strain that was used to generate display phages in this study.

| Prophage Name | Prophage Genotype | Source |
| --- | --- | --- |
| YXTL1026 | $\lambda cI857$, *Sam7*, $\Delta fhuA$, $\Delta b2$, *E<sub>E158K</sub>* | This study |
| YXTL1048 | YXTL1026 *D* ~linker:: *tetA-sacB* *,<br>The *tetA-sacB* cassette is inserted with a linker coding sequence at $\lambda D$ gene TAA stop | This study |
| YXTL1059 | YXTL1026<br>*tetA-sacB*::linker~*D*,<br>The *tetA-sacB* cassette is inserted with a linker coding sequence at $\lambda D$ gene ATG start | This study |
| YXTL1226 | XTL1026 *stf386-tetA-sacB* | This study |
| YXTL1439 | XTL1226 *stf386* truncation. *TetA-sacB* is replaced by sequence to truncate *stf*. | This study |
| YXTL1441 | XTL1439 *D*~linker::*tetA-sacB*, the *tetA-sacB* cassette is inserted with a linker coding sequence at $\lambda D$ gene ATG start. Used to create melanoma C-terminal $\lambda D$ fusion phage in a strain that included *Stf* truncation. | This study |
| YXTL1442 | XTL1439 *tetA-sacB*::linker~*D*, the *tetA-sacB* cassette is inserted with a linker coding sequence at $\lambda D$ gene ATG start. Used to create melanoma N-terminal $\lambda D$ fusion phage in a strain that included *Stf* truncation. | This study |

\* ~ symbol refers to a genetic fusion.

**Table 4C:**
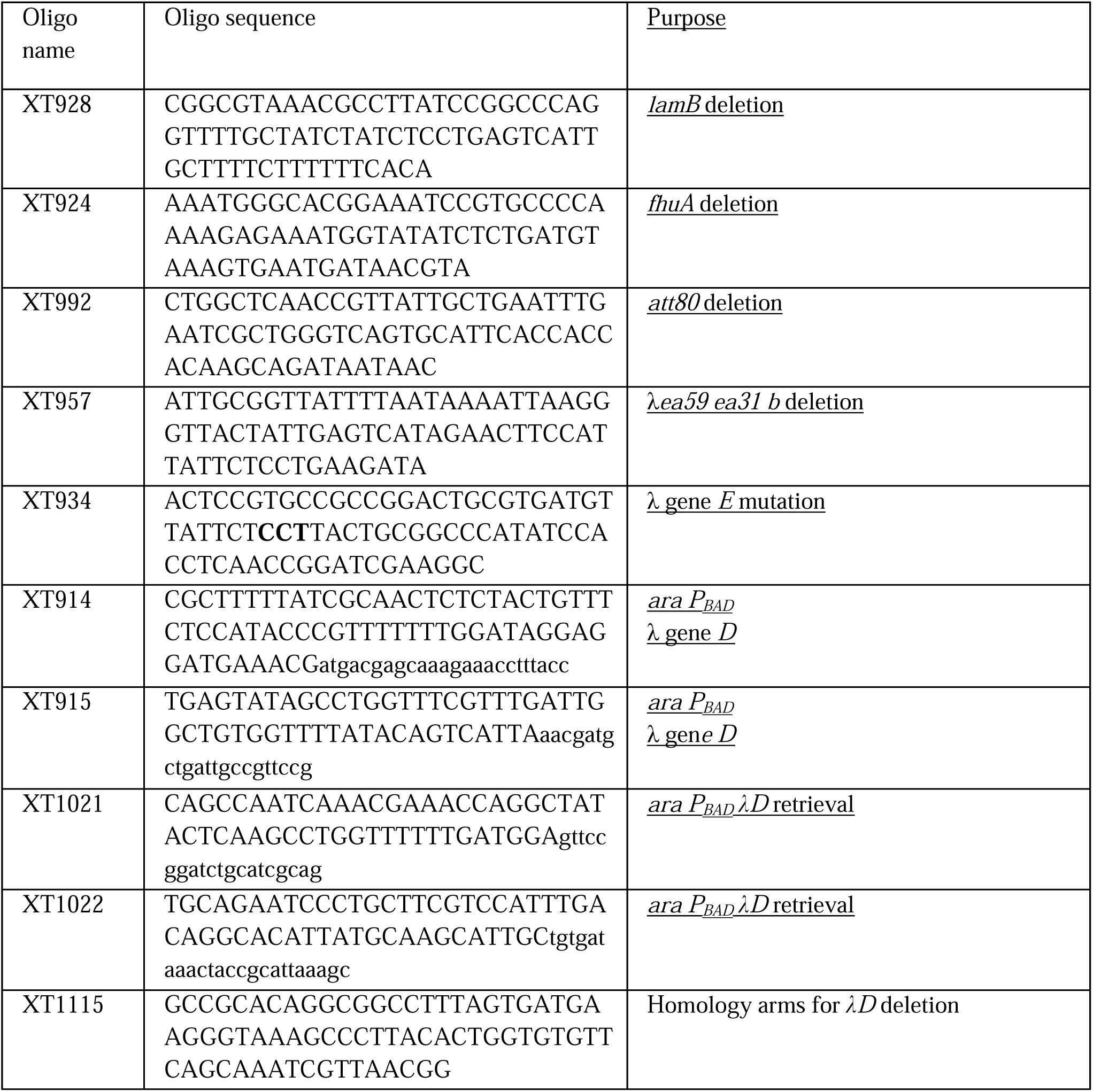
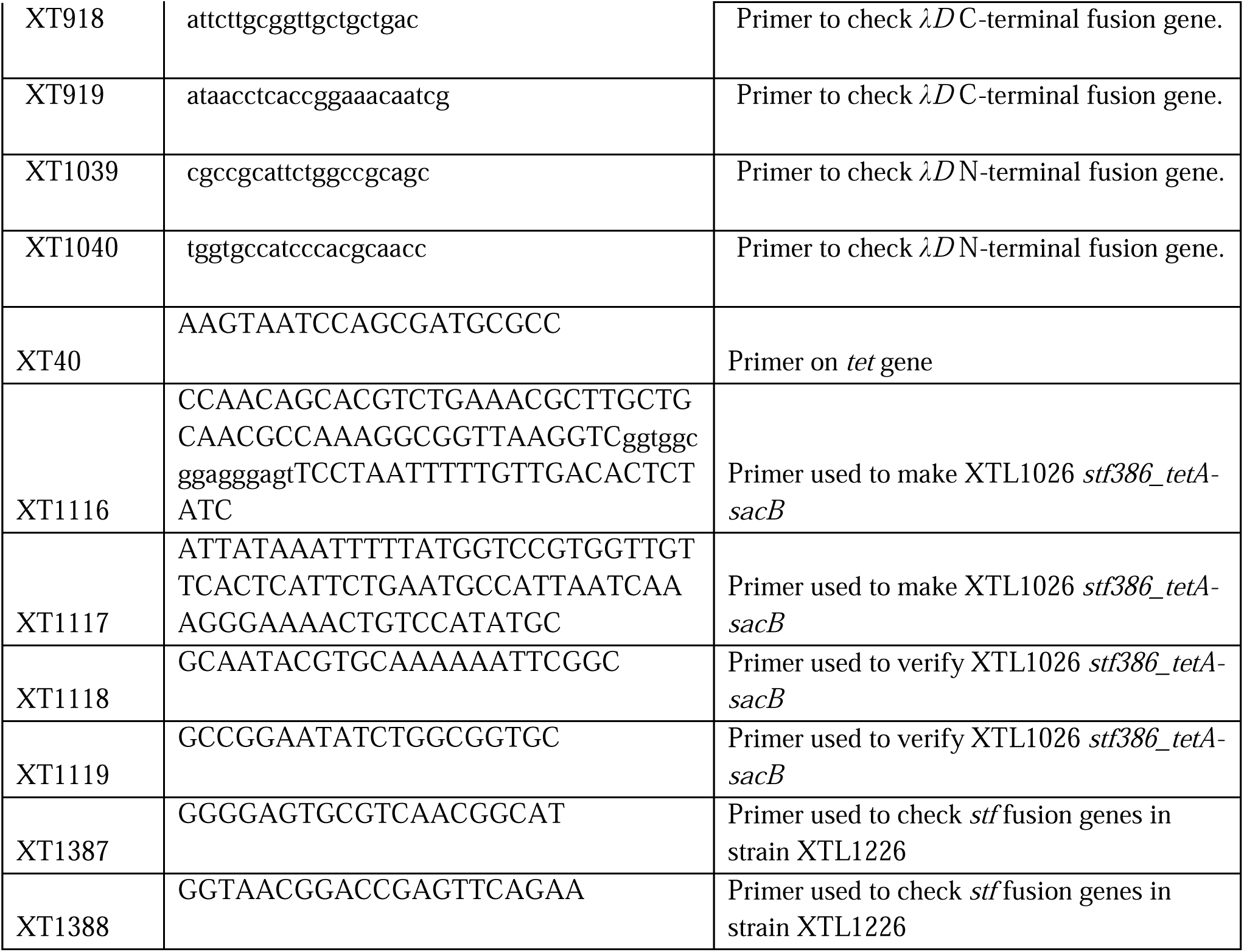
Oligonucleotides used to generate bacterial and prophage strains in this study. Designed for the deletion, modification, and addition of genes or to check by PCR and Sanger Sequencing the identity of modified sequences.

### *E. coli* strain evolution

#### The details of strain construction and constructs are adapted from Thomason et al., 2023^43^

*<u>SJ_</u>*<u>ΧΤ*L175*</u>: Strain SJ_XTL175 was used as the starting strain for the construction of the host bacterium used here. SJ_XTL175 is a wild-type MG1655 derivative^44^ that contains the following genotype: Δ*lacI, lacY*_A177C_, Δ*araE,* Δ*araFGH*::*spec^R^*, Δ*araD*::*tetA*-*sacB*-*amp*. SJ_XTL175 was further modified by recombineering to replace the arabinose operon genes *araB araA araD* with gene *D* of λ. The *araC* gene remains intact, allowing *AraC* regulation of the λ*D* gene (Fig. 6). In this construct, *tetA*-*sacB*-*amp* DNA is inserted after *araA* to create a counter-selection mechanism (Fig. 6B). Then, from the start codon of *araB*, including *araA* and the insert, to the *araD* TAA stop codon is replaced with gene *D* of λ (Fig. 6C). This places *D* of λ under *AraC* regulation and arabinose-inducible control. The other mutations (Δ*lacI, lacY*_A177C_, Δ*araE,* Δ*araFGH*::specR) already present in SJ_XTL175 confer on this new strain (XTL981) the ability to tune the λD-protein expression level in every cell in response to the added extracellular arabinose concentration^44^. A mixture of wild-type λD and fusion λD has been used before to stabilize λ display particles^45^, but it has not been used previously in a well-controlled system like the tunable arabinose method used here. In our system, wild-type λD protein expression can be turned on at a precise time and directly tuned to the concentration of arabinose added. This allows for a stable, high-titer phage yield from each lysis, which is especially necessary to support the stability of the phage particle when fusing larger display proteins^46^.

*<u>XTL981:</u>* Strain XTL981 described above has been further engineered to optimize λ phage production and the generation of λ particles carrying D-fusion proteins. First, XTL981 was infected with λ*cI857 Sam7* phage to generate lysogens, and standard techniques were used to identify a lysogen in which a single phage genome was integrated into its native attachment site (*attB*) on the bacterial chromosome^47^. Next, additional changes were made to the bacterial chromosome and to the prophage λ*cI857 Sam7* genome using recombineering technologies. As mentioned above, the host *lamB* gene was deleted. The *fhuA* gene, encoding the phage T1 and phage φ80 receptor protein, was deleted to avoid contamination by T1 and φ80 bacteriophage^48^. The nonessential λ genes *ea59* and *ea31* were precisely removed in a DNA segment of 4,285bp from the *b* region of the λ prophage^49^. This deletion provides more room in the phage genome for cloning and encapsidation of DNA segments to be fused to the *D* gene.

*<u>XTL1026:</u>* XTL1026 is the strain containing all the modifications to the host and the λ prophage that are described above. This strain also contains plasmid pKM208, which is a low-copy pSC101-based plasmid that expresses Red functions Exo Bet and Gam from the *lac* promoter for recombineering^50^. The *D* gene in the λ prophage is the target for creating *D* fusions to express conjugated proteins joined at either the N or C terminus of λD. Two strains are derived from XTL1026 in which the λ*D* gene is modified for generating λ*D* gene fusions. The counter-selectable *tetA-sacB* marker cassette is inserted just upstream or downstream of the *D* gene in the prophage by selecting for tetracycline resistance. These two Tet^R^ prophage strains are respectively, XTL1059 and XTL1048. Both become sucrose sensitive because of the *sacB* gene function linked to the *tetA* marker, which can then be replaced with the sequence of the fusion peptide. Strain XTL1048 was used to create the C-terminal SARS-CoV-2 spike protein λD fusion phage described in the results.

*<u>XTL1226:</u>* This strain was created from XTL1026 to allow for the addition of foreign peptides to the C terminus of λ s Stf protein. Initially, PCR was performed on the T-SACK strain^41^ using primers XT1116 and XT1117 (Table 4C) to generate the *stf386-tetA-sacB* cassette. The recombination of this cassette into the prophage strain inserts the *tetA-sacB* cassette into the *stf* gene at position 386 and can be verified by PCR using primers XT1118 and XT40 (expected product: 505 bp). Strain XTL1226 is sucrose sensitive due to the addition of the *stf386-tetA-sacB* cassette.

This strain is then able to recombine with foreign peptides. When replacing the *stf386-tetA-sacB* cassette with a foreign peptide, we use a gene block that truncates the rest of the *stf* gene from its 774 amino acid wild-type length to 386 amino acids, placing the fusion peptide at the new *stf* C terminus. This truncation slightly improves phage titer and was performed to ensure fusion peptide stability to a smaller Stf fusion protein. Full-length Stf is non-essential to phage assembly, lysis, or protein display, so it can be truncated for optimal fusion without disrupting functions necessary to protein display. XTL1226 was used to produce Stf melanoma epitope prophage by the insertion of melanoma epitopes with flaked λ homology at the *stf386-tetA-sacB* cassette location (Table 1 and Table 2). The prophages were screened using primers XT1118 and XT1119 (expected product: 436 bp). Identity of melanoma epitopes were further confirmed by Sanger sequencing.

*<u>XTL1439</u>:* This strain completes the truncation of Stf without placing a fusion peptide at the Stf location. Strain XTL1226 is recombineered with a gene block to replace the *stf386-tetA-sacB* cassette with a stop codon, truncating Stf. A new *tetA*-*sacB* cassette is then inserted at the C or N terminus of this strain’s λ*D* gene to create strains XTL1441 and XTL1442. These two strains were used to then make melanoma epitope λD display phages in a strain with a truncated Stf to create an identical genetic background for comparison with the Stf melanoma epitope display phages.

### Prophage induction procedure

Once foreign proteins have been genetically fused to the *D* or *stf* gene of the λ lysogen, the prophage can be induced to create a bacterial lysate which may be further purified and used in animal studies. To induce prophage, the *E. coli* strain containing the fusion prophage is grown at 32° C in LB broth to 2×10^8^ cell/ml before quickly shifting to 42° C for 30 min and back to 39° C for 3 hrs (Fig. 7). Additionally, 0.2% total arabinose can be added to induce host chromosome wild-type λD to mitigate potential phage particle instability caused by the fusion (Fig. 6). Due to the *Sam7* mutation that inactivates the phage holin protein, a lysate of the induced phage can be prepared by at least two methods. The first is the addition of chloroform, and the second is spinning the cells down into a pellet, resuspending in a small volume, and freezing overnight at –80° C. After freezing, the resuspended pellets are thawed to lyse the cells. The resulting crude lysate is then spun down to form a pellet and the supernatant filtered through 0.45 and 0.2 micron filters to collect the clear phage lysate. Freeze/thaw was the method used for lysis in the studies described here. Whether made through freeze/thaw or chloroform addition, the lysate is then titered on a *supF* strain (XTL1212) which suppresses the *Sam7* mutation to make gpS and allow lysis and plaque formation. Strain XTL1212 also contains the plasmid pLXT42, which produces λD protein to complement D-fusion phage particles that may be somewhat defective in D function and cannot form plaques efficiently on their own. The λD fusion identity can then be confirmed by western blot and DNA sequencing before the phage lysates are purified by tangential flow filtration.

**Fig. 7.**
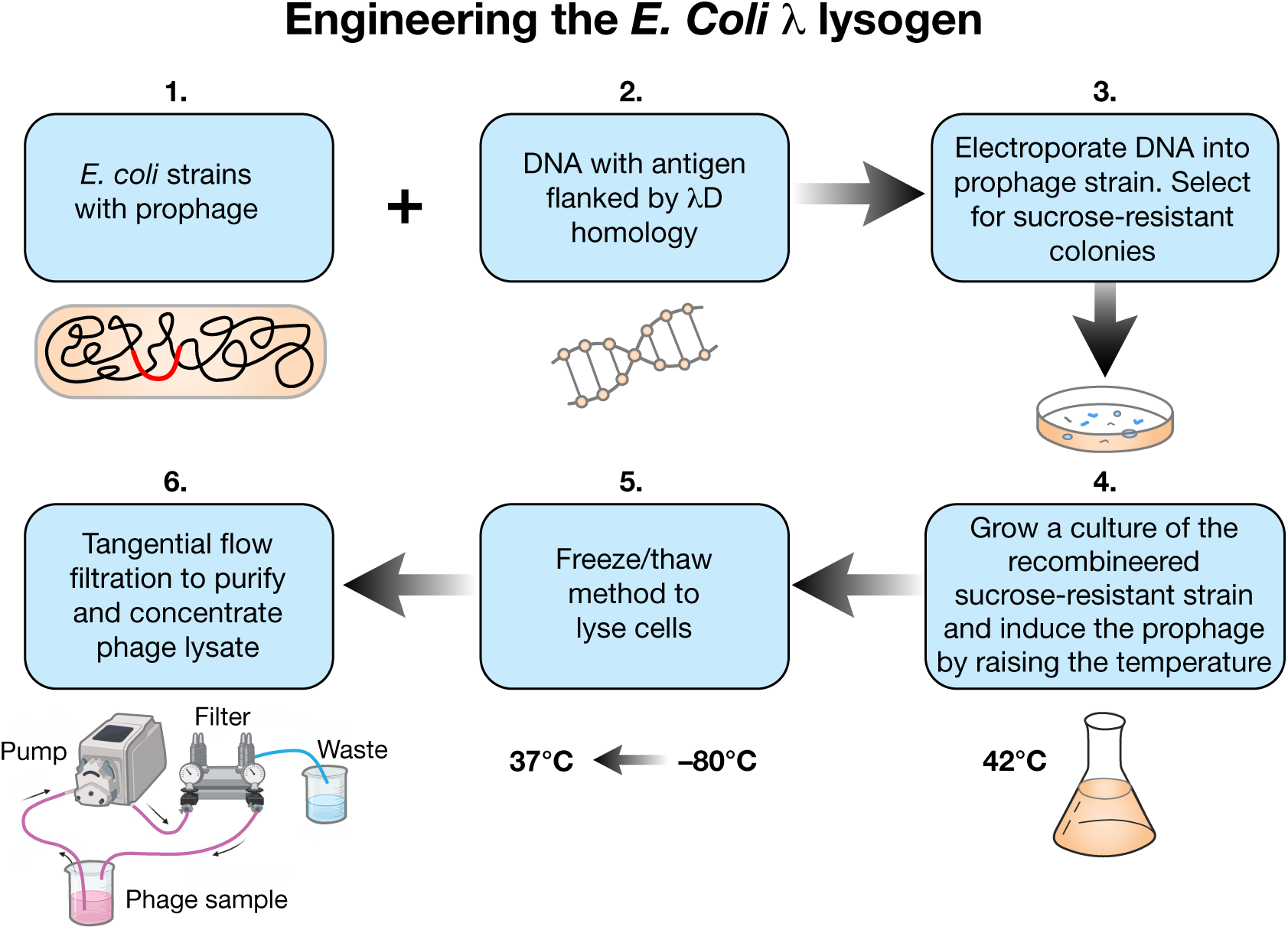
Process of prophage engineering and phage lysate production. 1. *E. coli* strains with prophage, 2. foreign DNA flanked by λD homology, 3. The DNA is electroporated into the prophage strain, and the strain is plated on sucrose media to select for recombinants. 4. Once recombinant colonies are confirmed, the selected strain is grown at 32° C in LB broth to 2×10^8^ cell/ ml. Prophages are induced by shifting the water bath temperature to 42° C for 30 minutes and then down to 39° C for another 3 hours. 5. Culture containing assembled phages within *E. coli* cells is spun down, resuspended in a small volume, frozen at –80° C overnight, and then thawed the next day at 37° C to lyse the culture by the freeze/thaw method. 6. Phage contained in the supernatant lysate from the freeze/thaw step are then purified and concentrated by tangential flow filtration (TFF) at room temperature. During TFF, a tube system feeds phage lysate through a filter that takes bacterial endotoxin to waste and feeds phage back to the starting sample. Running the phage through this process many times purifies out endotoxin and concentrates purified phage in a small volume.

### Tangential flow filtration

Engineered phage lysates were filtered and their titer concentrated by passage through a Pellicon® Mini Cassette 0.1 mm Composite Regenerated Cellulose filter by Millipore. The filter retains the phage particles and allows bacterial waste to pass through and be discarded. Filtration occurs by running 5-7 liters of 10 mM Tris and 50mM MgCl□ buffer^33^ through the filter system to purify phage. Phage are resuspended after filtration in a small volume (5-15 ml) of the same buffer. The MgCl□ stabilizes phage particles and allows for long-term storage. This buffer can be diluted in PBS before injection into mice to reduce MgCl□. Endotoxin readings were taken after filtration using the Endosafe® Nextgen-PTS™ (Portable Test System) by Charles River to ensure that the lysate was below an acceptable level of toxin for animal injection.

### Mouse study protocol

The mouse study conducted to test the immunogenicity of XTL1382 display phage utilized six– to eight-week-old Balb/c female mice. Groups of five mice were injected with antigen fusion phage, vector phage, and 10 mM Tris and 10mM MgCl□ buffer. Mice were injected intraperitoneally with 500 µl doses of 2×10□ pfu three times over the course of the 49-day study, on days 0, 13, and 34. Bleeds were taken at five time points on days 0, 13, 27, 34, and terminally on day 49. This study was performed under Integrated Biotherapeutics’ (Rockville, MD) approved IACUC protocol #160805.

### Analysis: ELISA and western blot

#### Western blot

##### 1. Western Blot identification of SARS-CoV-2 spike protein epitope using SARS-CoV-2 infected and healthy donor human serum

To identify antibody-protein affinity between spike antigen display phage and human serum samples, we ran phage samples on 4-20% SDS gradient gels. The samples were then transferred to a nitrocellulose membrane for standard western blotting. First the membrane was blocked with 2.5% non-fat milk buffer, then it was triple washed with TNE buffer (10mM Tris-HCl, 50mM NaCl, 2.5mM EDTA) prior to incubating with 1000-fold dilution of human serum samples extracted either from a healthy donor or a patient positive for SARS-CoV-2. The membrane was incubated with serum samples for three hours and then washed with a wash buffer (10mM Tris-HCl, 50mM NaCl, 2.5mM EDTA). It was then incubated with 10,000-fold diluted goat anti-human IgG secondary-HRP conjugated antibody for an hour and washed with a wash buffer containing 0.1% tween. The membrane was developed using a clarity enhanced chemiluminescence substrate from Bio-Rad.

##### 2. Western blot protocol for mouse serum antibody affinity

Western blots were conducted to quantify mouse sera antibody affinity to phage sample fusion protein. They were performed by loading fusion phage and vector phage on an SDS 4-20% tris-glycine gel, transferring separated proteins to a nitrocellulose membrane, blocking with 5% milk buffer in 1X TNE, and probing with 1:1000 dilution of mouse serum in milk buffer, followed by 1:10,000-fold goat anti-mouse IgG secondary-HRP in 1X TNE. Membranes were developed with Clarity Max Western ECL substrate and imaged with chemiluminescence.

#### ELISA

*ELISA Assay protocol adapted from Hong Zhou (NIH/NCI) and Thermo Fisher’s general ELISA protocol:* (https://www.thermofisher.com/document-connect/document-connect.html?url= https://assets.thermofisher.com/TFS-Assets%2FLCD%2Fmanuals%2FD20692∼.pdf)

Mouse serum was analyzed for the presence of antibodies specific to the injected λD XTL1382 peptide by ELISA (enzyme-linked immunosorbent assay). ELISA demonstrated antibody-protein binding by incubating a synthetic peptide version of the fusion phage peptide on a nunc-immobilon plate at a 20 µg/ml concentration. Once the synthetic peptide was immobilized on the plate, mouse serum was added in several 10-fold dilutions. The synthetic protein/antibody binding interaction was quantified by adding TMB substrate, stopping the reaction after 12 min with 17% H□SO□ stop solution, and reading color intensity at 450 nm. Absorbance was read at a 100x dilution. After subtracting the background, collected at 562 nm, data was plotted in GraphPad Prism 10. Error bars were generated in Prism software.

## Materials

### Synthetic peptides created by GenScript Inc

● Spike Protein Epitope XTL1382: YDPLQPELDSFKEELDKYFKNHTSPDVDLGD Synthetic peptide was custom synthesized by GenScript Inc. Peptide sequence is from the second subunit of the SARS-CoV-2 spike protein. The purity of this peptide was above 95%. It was resuspended in sterile H O for use in ELISA at 20 µg/ml in each well for the detection of antibodies.

### Antibodies

● Anti-λD antibody: Custom-made polyclonal anti-λD capsid protein antibody raised in mouse against purified λD protein from GenScript Corp.
● Human serum used in antigen selection: blood serum samples from three healthy donors and three patients positive for SARS-CoV-2 infection were generously gifted by the laboratory of Robert J. Kreitman at NCI/NIH.
  ○ Healthy Donors:
    ▪ α-COVI-SP-0005
    ▪ α-COVI-SP-0007
    ▪ α-COVI-SP-0008
  ○ COVID+ Sera
    ▪ α-COVI-SP-0009
    ▪ α-COVI-SP-0179
    ▪ α-COVI-SP-0187

## Notes

### Competing Interest Statement

The authors have declared no competing interest.

