## Supplemental Figure 1 for "A λ Phage Platform for Successful Therapeutic Display of Protein Antigens"

A.

**Mouse Study Timeline**

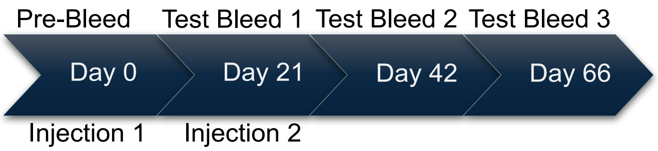


B.

**XTL1382 Sera Antibody ELISA Titer**

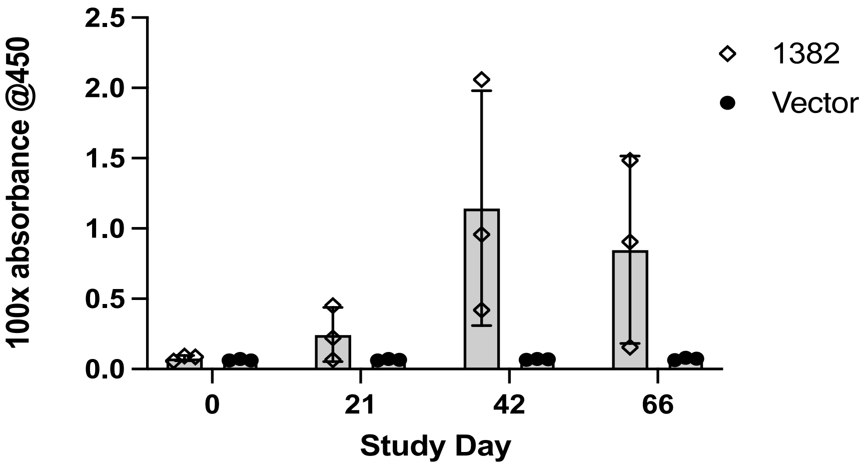


C.

**Phage Sample Western Blot Against Phage-Injected Day 42 Sera**

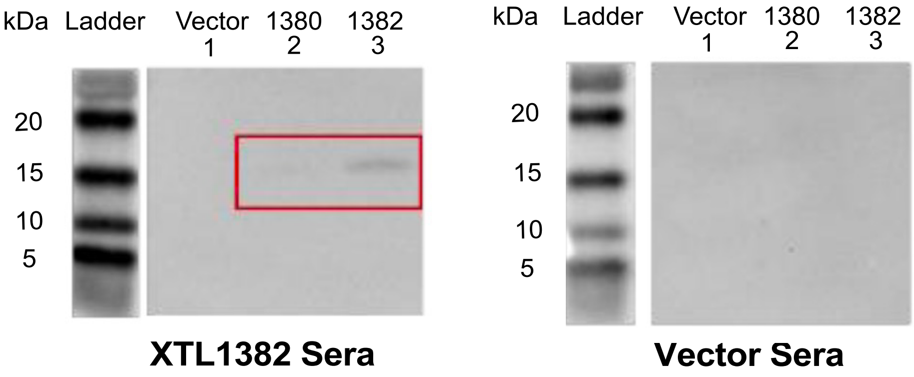

**Fig. S1. Preliminary antibody generation study with vector λ phage (XTL1026) and λ phage displaying SARS-CoV-2 spike protein epitope XTL1382.** A: Timeline of a preliminary mouse immunogenicity study. Vector and XTL1382 fusion phage were injected in groups of three mice via an IP route on Days 0 and 21 with 500 µl doses of 1x109 pfu. Sera was gathered on Days 0, 21, 42, and 66. B: ELISA antibody titer absorbance data at 100x dilution, testing vector and fusion phage mouse sera groups (n=3) against the SARS-CoV-2 synthetic XTL1382 peptide. Antibody titer was measured at each time point over the course of the study with a single reading per mouse and three mice per bar. Absorbance indicates a varied but strong response from the XTL1382 experimental group sera, especially on Day 42 and 66. Vector-injected mice did not demonstrate antibody signal against peptide XTL1382. C: Western blot of vector, XTL1380, and XTL1382 display phage blotted against Day 42 XTL1382-injection serum (left) and Day 42 vector-injection serum (right). When blotted against XTL1382-injection serum, the XTL1382 phage lane (3) produced a band. The same phage blotted against vector-injection serum did not produce a band. These results indicate that an injection of XTL1382 spike protein epitope displayed on phage stimulated an antibody response specific to the spike protein XTL1382 epitope.
